# A tissue-intrinsic mechanism sensitizes HIV-1 particles for TLR-triggered innate immune responses

**DOI:** 10.1101/2023.11.22.568261

**Authors:** Samy Sid Ahmed, Liv Zimmermann, Andrea Imle, Katrin Wuebben, Nadine Tibroni, Lena Rauch-Wirth, Jan Münch, Petr Chlanda, Frederik Graw, Oliver T. Fackler

## Abstract

In vivo, HIV-1 replicates within tissue, but how three-dimensional (3D) tissue-like environments influence viral spread and pathogenesis remains largely unknown. We previously identified an *Environmental Restriction to cell-free Virus Infectivity* (ERVI), imposed by collagen-rich 3D extracellular matrix (ECM), which diminishes HIV-1 particle infectivity. Here we report that ERVI is broadly implemented by various adhesive ECM components assembled into tissue-like 3D scaffolds. This restriction rapidly reduces infectivity within minutes, is saturable, and affects diverse primary HIV-1 strains and virions with distinct viral glycoproteins by impairing their ability to fuse with target cells. Importantly, particles subjected to ERVI also trigger pronounced pro-inflammatory cytokine secretion by monocyte-derived macrophages. Mechanistic analyses reveal that transient contact with collagen fibers induces conformational changes in the viral envelope glycoprotein (Env) and enhance its recognition by toll-like receptor (TLR) 2. This recognition promotes routing of viral particles into TLR8-positive endosomes, amplifying innate immune sensing of viral RNA genomes. ERVI thus acts via a dual mechanism: directly limiting the fusogenicity of cell-free virions and sensitizing them for innate immune detection. These findings highlight the biophysical properties of the ECM as a previously unrecognized, tissue-intrinsic barrier that restricts HIV-1 spread and promotes local inflammation representing a novel, broadly acting arm of antiviral innate immunity.

## Introduction

Untreated HIV infection induces a complex pathology characterized by depletion of CD4 T cells, generalized dysfunction of B and CD8 T cells as well as chronic immune activation and inflammation that promotes lymph node fibrosis^1, 2, 3^. In particular immune cell dysfunction and chronic inflammation persist even in people living with HIV (PLWH) in which peripheral viral load is suppressed by highly active antiviral therapy^4^. While extensive research over the past decades provided a detailed understanding of the replication strategies and molecular interactions that enable HIV to replicate in individual target cell types *ex vivo*, much less information is available regarding the mechanisms of HIV-induced immunopathology in infected tissue. This lack of knowledge on the impact of tissue on HIV spread and pathogenesis reflects the limited availability of culture models to address these questions experimentally. Humanized mouse models provide a valuable system to study virus spread *in vivo* but fail to recapitulate important immune responses of PLWH and their long-term effects. Organotypic systems such as tonsil or cervix explants allow to study HIV infection in the context of physiological tissue composition but experimental parameters are difficult to control ^5, 6^. We previously established three dimensional (3D) matrices of the major extracellular matrix (ECM) component type I collagen as a cell culture model for extracellular tissue environment in which parameters such as cell density and 3D organization can be controlled to complete the portfolio of experimental systems to study HIV spread ^5, 7^. The initial characterization of this 3D collagen culture model allowed us to address which transmission mode HIV employs in tissue-like environments: HIV can spread from infected donor cells either by the release of cell-free particles into the extracellular space that then diffuse to infect new target cells (cell-free transmission) or via close physical contacts between donor and target cell (cell-associated transmission)^8, 9, 10^. Combining computational modelling and subsequent experimental validation identified cell-associated transmission as the predominant mode of virus spread in 3D collagen at conditions of limited cell density where virions either need to diffuse or donor cells have to migrate to new target cells to sustain HIV spread^7^. In contrast, in 3D collagen with very high cell density or in classical suspension cultures, cell-free and cell-associated transmission modes supported HIV-1 spread with comparable efficacy. This shift towards cell-associated virus transmission in tissue-like environments reflected an enhanced efficiency of cell-associated spread, but also a significant reduction in the infectivity of HIV-1 particles. Imaging analysis revealed that HIV-1 particles freely diffuse in the tissue-like environment but undergo short (milliseconds) and reversible physical interactions with collagen fibers. These findings suggested that in tissue, the physical contact with ECM limits the infectivity of cell-free HIV-1 particles but the mechanism of this restriction remained elusive. As an enveloped virus particle, the infectious potential of HIV-1 virions is determined by a large number of parameters. This comprises basic properties such as particle integrity, packaging of the viral genome and essential enzymes, incorporation of the viral glycoprotein Env, and liquid-order membrane microdomain-like lipid composition of the viral envelope^11, 12, 13, 14^. Dependency on this complex set of requirements renders the generation of infectious HIV particle an attractive target of the activity of cell intrinsic antiviral factors (so called restriction factors) including tetherin, SERINC proteins, 90K, IFTIM proteins, GBP 2 and 5 and PSGL-1 that reduce HIV-1 infectivity via distinct mechanisms^15, 16, 17, 18, 19, 20, 21, 22^. In analogy to these intracellular barriers that counteract HIV-1 spread, we designated the decrease of virion infectivity by tissue-like environments as an *Extracellular Restriction of Virion Infectivity* (ERVI).

HIV particles also present a number of pathogen-associated patterns (PAMPs) that can be recognized by pattern recognition receptors (PRRs) to trigger antiviral signaling. HIV-associated PAMPs include the incoming single strand RNA genome (gRNA)^23^ and products of reverse transcription of the RNA genome into DNA but also post integration replication intermediates^24^ and protein structures^25, 26, 27^. Whether tissue environments impact the mode and potency of innate immune recognition of HIV-1 particles has not been assessed.

In this study, we set out to define the mechanism and functional consequences of ERVI. We find that ERVI has two mechanistically distinct effects on HIV-1 virions that (i) restrict their infectivity and (ii) sensitizes them for innate immune recognition. Mechanistically, the infectivity impairment results from reduced fusogenicity of virions while enhanced innate immune recognition reflects the recognition of the viral glycoprotein by TLR-2 and sensing of viral gRNA by endosomal TLR-8. These results uncover that tissue-like environments bear a previously uncharacterized intrinsic antiviral activity that limits viral spread while inducing an innate immune response in myeloid cells.

## Results

### ERVI is a rapidly induced, saturable and conserved restriction to HIV-1 infectivity

Our previous results had established that tissue-like 3D collagen environments pose the ERVI barrier to markedly reduce the infectivity of HIV-1 particles recovered from the supernatants of 3D collagen matrices in which virus producing cells or cell-free virus particles had been embedded^7^. Single particle tracking revealed that HIV virions diffuse freely in the 3D matrix but undergo transient physical contacts with collagen fibers. These interactions were in the millisecond range and did not result in coating of the fibers with virus particles^7^. To gain more insight into the nature of ERVI, we compared the impact of culturing HIV-1 NL4.3 particles in suspension or upon embedding in 3D collagen with different densities over time (dense collagen (DC): 3 mg/ml rat tail collagen; loose collagen (LC): 1.6 mg/ml bovine skin collagen) (Fig. 1a). The infectivity of virions that had diffused into the culture supernatant was assessed on TZM-bl reporter cells kept in the absence of collagen, in which the luciferase gene is under control of the HIV-1 promoter and *de novo* expression of the viral transactivator Tat in productively infected cells triggers luciferase expression. The amount of luciferase expression relative to the amounts of virus used for infection, as determined by quantifying the activity of viral reverse transcriptase by the SG-PERT assay, yielded the relative infectivity of HIV particles. In line with our previous findings, single rounds of infection revealed that culturing HIV-1 in DC or LC for 16h reduced their infectivity to 10.6% or 14.3% of the particles from parallel suspension cultures (Fig. 1b). Kinetic analysis revealed that virion infectivity was reduced immediately or within 4h after contact with 3D collagen in DC and LC, respectively. The subsequent reduction of the remaining virion infectivity over time followed comparable kinetics under all culture conditions (Fig. 1c). A significant reduction of virion infectivity was also observed when virions were cultured on top of already polymerized collagen matrices (Fig. 1d, see Sup. Fig. 1a for experiment layout: residual infectivity for viruses embedded in collagen: 6.6% DC, 32.4% LC or viruses kept on top of collagen: 26.7% DC, 43.5% LC). In contrast, supernatants of collagen matrices polymerized in the absence of virus particles or the buffers in which collagen is polymerized (PA) failed to restrict the infectivity of virions. The reduction of virion infectivity by collagen matrices thus depends on physical contact of virions with the matrix and is not mediated by soluble components. Physical stress can impair virus infectivity by inducing shedding of the viral glycoprotein from the virion, and the efficacy of glycoprotein incorporation is an important determinant of infectivity^28, 29^. However, HIV-1 Env gp120 levels were comparable between virions cultured in suspension or in 3D collagen (Fig. 1e,f). Since cellular restrictions to virus infection typically act as physical barriers that can be overcome by an excess of virus particles^30^, we analyzed if the amounts of virions added to a constant culture volume affected the magnitude of reduction in virion infectivity (Fig. 1g). While the relative infectivity of virus particles in suspension was unaffected by the concentration of virus, the infectivity reduction by ERVI was significantly less efficient at higher virus concentrations in DC or LC (Fig. 1g; relative infectivity 7- or 2-fold higher at 10^7^ vs. 10^5^ infectious units in DC or LC; p<0.09 and p<0.015 respectively). The infectivity restriction by ERVI is therefore saturable.

**Figure 1:**
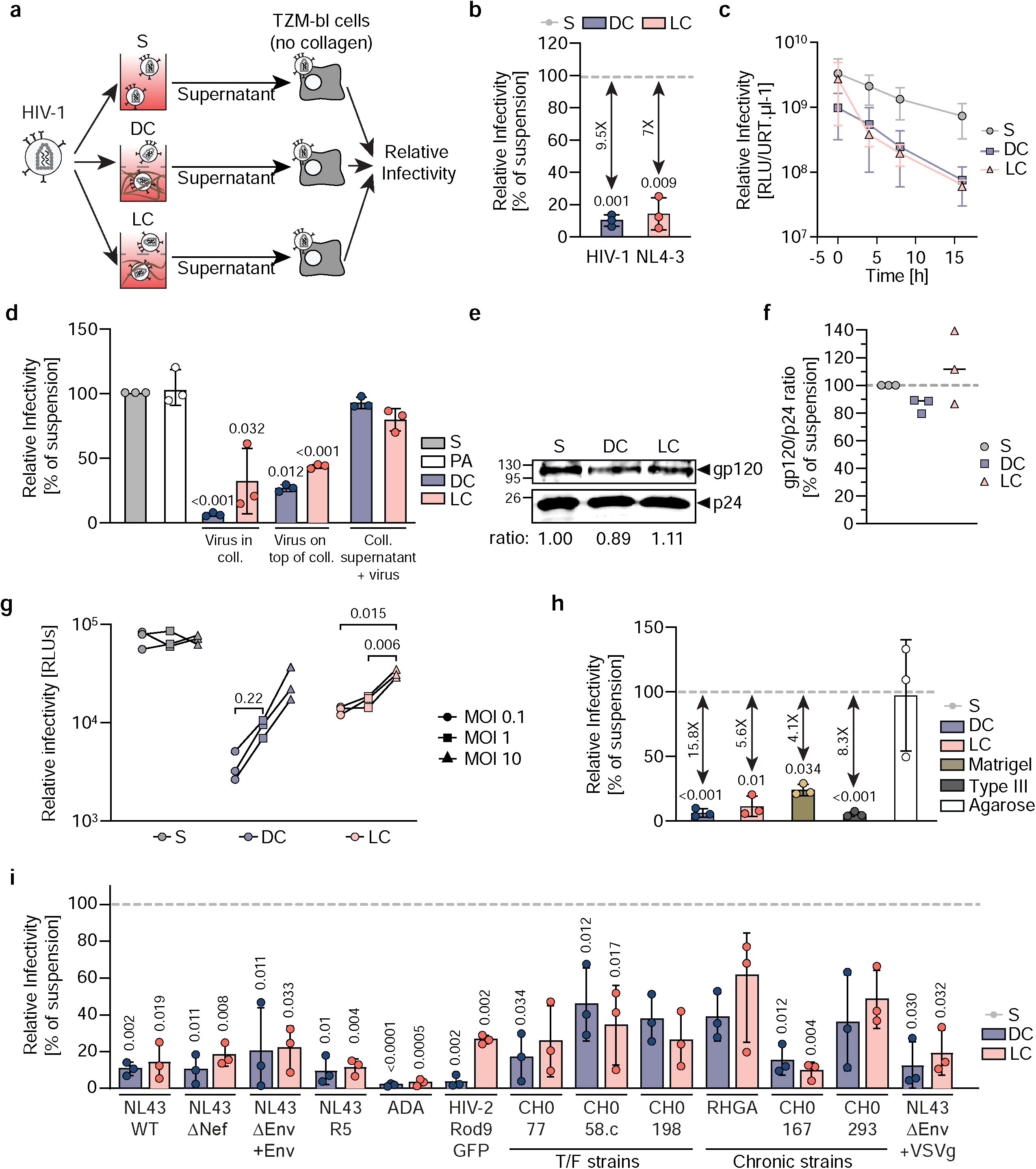
ERVI is Rapidly Induced, saturable, and conserved across viruses and different types of matrices. **a**. Experimental workflow. HIV-1 NL4.3 virions were cultured in suspension (S) or embedded in collagen matrices (DC: Dense collagen; LC: Loose collagen). TZM-bl cells were infected and the relative infectivity determined. **b**. Relative infectivity of HIV-1 NL4.3 particles. Virions were cultured in S, DC or LC for 16h prior to relative infectivity determination. Data is normalized to suspension condition (grey dotted line). **c**. Kinetics of ERVI. Culture supernatants were harvested at 0, 4, 8 or 16h post seeding/embedding, the relative infectivity of virions was determined as in b. **d**. Relative infectivity of differentially cultured HIV-1 NL4.3 virions. PA: polymerizing agent. **e**. Western blot analysis showing the gp120/p24 ratio from HIV-1 NL4.3 virions cultured in S, DC or LC. The gp120/p24 ratios are indicated below the blots. **f**. Quantification of western blot analysis as in e. **g**. Saturation of ERVI. Increasing amounts of virions were cultured in S, DC or LC. Their relative infectivity was determined as in b. **h**. Relative infectivity of HIV-1 NL4.3 particles after seeding or embedding in different matrices for 16h. Data is normalized to suspension condition (grey dotted line). **i**. Conservation of sensitivity to ERVI across lentiviruses. The relative infectivity of a panel of lentiviruses was assessed after ERVI as in a. Data is normalized to respective suspension condition (grey dotted line). Results represent the mean ± SD of 3 independent experiments. Symbols indicate data from individual experiments. Significance is indicated by p-values, and was calculated by one-way ANOVA test; Dunnett’s post-test (**b,d,h,i**), or by matched two-way ANOVA test; Tukey’s post-test (**g**).

We next sought to define how the architecture and biophysical properties of the 3D matrix affect its ability to reduce the infectivity of HIV-1 virions and compared several matrices that (i) can be polymerized without harming *per se* the infectivity of HIV particles (e.g. high temperature or UV exposure) and (ii) result in pore sizes that allow HIV-1 particles to diffuse within and out of the 3D matrix (Fig. 1h, Sup. Fig. 1b). DC and LC matrices are assembled into fibers polymerized from purified type I collagen proteins, the most abundant collagen type in tissue^31^. Confocal reflection microscopy analysis of these matrices confirmed the different density of both densities of type I collagens after polymerization and revealed that DC was enriched in branched collagen bundles (Sup. Fig. 1b). The architecture of 3D matrices made of type III collagen, the second most abundant fibrillary collagen in tissue that is synthesized by reticular cells and lines e.g. vasculature^32^, resembled that of DC, albeit with shorter and thinner collagen bundles, and type III collagen reduced HIV-1 infectivity with similar efficacy than type I collagen. We also tested Matrigel, a complex extracellular environment of the basal membrane rich in collagen IV, III and I^32^ as model for a complex tissue environment. Although assembles into 3D matrices with smaller pore sizes, refractive indexes and higher heterogeneity than purified collagen matrices which reduces signals from autoreflection microscopy (Fig. S1b)^33^, its antiviral activity was comparable to that of DC (24.1+/- 4.4% of suspension or 4-fold reduction). In contrast, agarose hydrogels, which are made of bundles of thin linear filaments without cell adhesion features that do not produce reflection signals^34^, did not impair the infectivity of HIV-1 virions. We conclude that ERVI is a conserved feature of adhesive collagen matrices and ECM.

To assess how conserved the sensitivity to ERVI is among primary lentiviruses, we next analyzed the impact of 3D collagen on a panel of lab-adapted and primary HIV-1 strains as well as one HIV-2 strain on TZM-bl reporter cells (Fig. 1i). The results revealed that the infectivity of all HIV strains tested was significantly reduced by ERVI, but to varying magnitude ranging from strong (52.9-fold, HIV-1 ADA, DC) to very mild inhibition (1.6-fold, HIV-1 RHGA, LC). The reduction of infectivity by ERVI was independent of the HIV-1 entry co-receptor preference but patient-derived transmitted founder and chronic HIV-1 variants tended to be less sensitive to ERVI than lab-adapted HIV-1 (DC: 10.5% residual infectivity for lab adapted vs 33.1% for primary isolates). Notably, also virions lacking HIV-1 Env but pseudotyped with the glycoprotein of vesicular stomatitis virus (VSVg) were sensitive to ERVI. Although different amounts of virus particles had to be used to achieve comparable infection rates for the different strains (see methods), the sensitivity of different virus strains to ERVI was not correlated to the amount of virus used for infection (Sup. Fig. 1c). Sensitivity to ERVI thus appears to be an intrinsic property of individual HIV variants. To specifically analyze the impact of the viral glycoprotein for sensitivity to ERVI, we pseudotyped GFP-encoding lentiviral particles with glycoproteins of several unrelated viruses. The transduction efficiency of lentiviral particles pseudotyped with the Hepatitis C Virus isolates Con1 or JFH1 or the Vesicular Stomatitis Virus glycoprotein (VSVg) was moderately reduced upon contact with 3D collagen (Sup Fig. 1f, statistically significant reduction for particles pseudotyped with VSVg glycoprotein in DC). Virus particles pseudotyped with other glycoproteins such as Influenza HA or the Ebola virus glycoproteins already lost all detectable infectivity within 16h of culture in suspension, precluding the analysis of the impact of 3D matrices (Sup. Fig. 1d,e). Together, these results suggest that the viral glycoprotein is a key determinant for the sensitivity to ERVI. Reduction of virus infectivity is observed with glycoproteins of different receptor specificity and topology but type I (HIV) and type III (VSVG) glycoproteins may be more sensitive to ERVI than type II glycoproteins (HCV).

Together, these results reveal that a broad range of HIV-1 variants are sensitive to ERVI. This extracellular restriction is exerted rapidly upon contact with the 3D environment, can be saturated by excess of virus particles, and is exerted by a variety of adhesive extracellular matrices.

### ERVI restricts HIV-1 infectivity without affecting structural integrity of virus particles

We next addressed in more detail how the physical contact of HIV particles with the 3D collagen environment affects their infectivity. We considered that following transient contacts of HIV-1 particles with collagen fibers, collagen material may remain attached to the virions and compromise their infectivity. To address this possibility we generated HIV-1 particles that incorporated a Vpr-integrase.GFP (Vpr.Int.GFP)^35^ fusion protein for visualization and remained sensitive to the infectivity reduction by ERVI (Fig 2a). Embedding these particles in fluorescently labelled LC allowed to assess the presence of fluorescent collagen at their surface by light microscopy (Fig 2b). Confocal imaging of virions within fluorescent LC matrices allowed to visualize a number of non-tethered particles present between collagen fibers (Fig. 2c). We next assessed a potential presence of fluorescent collagen fibers at the surface of Vpr.Int.GFP containing virions by TIRF-microscopy (Fig. 2d). While small fragments of free collagen fibers could occasionally be observed, we did not detect any AlexaFluor-647 signals studding the surface of virions (Fig. 2d). To analyze the impact of 3D collagen on virion morphology by cryo-electron tomography, HIV-1 particles kept in suspension or in dense or loose 3D collagen were placed on EM grids and processed for cryo-ET analysis. Under all three conditions, enveloped HIV-1 particles with the typical conical core and a diameter ranging from 104 to 154 nm (mean values: suspension: 134.5+/- 11.5 nm, dense: 135.2 +/- 9.6nm, loose: 132.0 +/- 11.6 nm) were observed (Fig. 2e,f). All analyzed particles appeared intact without appreciable membrane rupture or deformation and showed sparsely distributed Env spikes. As size comparison, we also analyzed the morphology of DC and LC fibers, which displayed the characteristic structure of collagen fibrils, with a tight packing of D-periodic polyproline type II helices corresponding to the spacing between individual tropocollagen monomers^36^ (Fig. 2g). Consistent with our analysis using fluorescent collagen, we failed to observe virion-associated collagen fibers (Fig. 2e). Since these analyses did not detect morphological aberrations in HIV-1 particle architecture resulting from the interaction with 3D collagen, we next asked if the reduced infectivity of HIV-1 particles subjected to ERVI can be rescued by infectivity-enhancing peptide nanofibrils (PNF) which boost the infectivity of HIV-1 particles by increasing their interaction with target cells and promoting viral fusogenicity ^37, 38, 39^. Indeed, incubating HIV-1 particles with EF-C or RM-8 PNFs increased the infectivity of all particles and almost fully overcame the inhibitory effect of ERVI (Figs. 2h,i). Together, these results reveal that ERVI does not result from global disruption of HIV-1 particle architecture and suggest that ERVI does not affect the intrinsic replication potential of HIV particles but rather the efficacy of their interaction with target cells.

**Figure 2:**
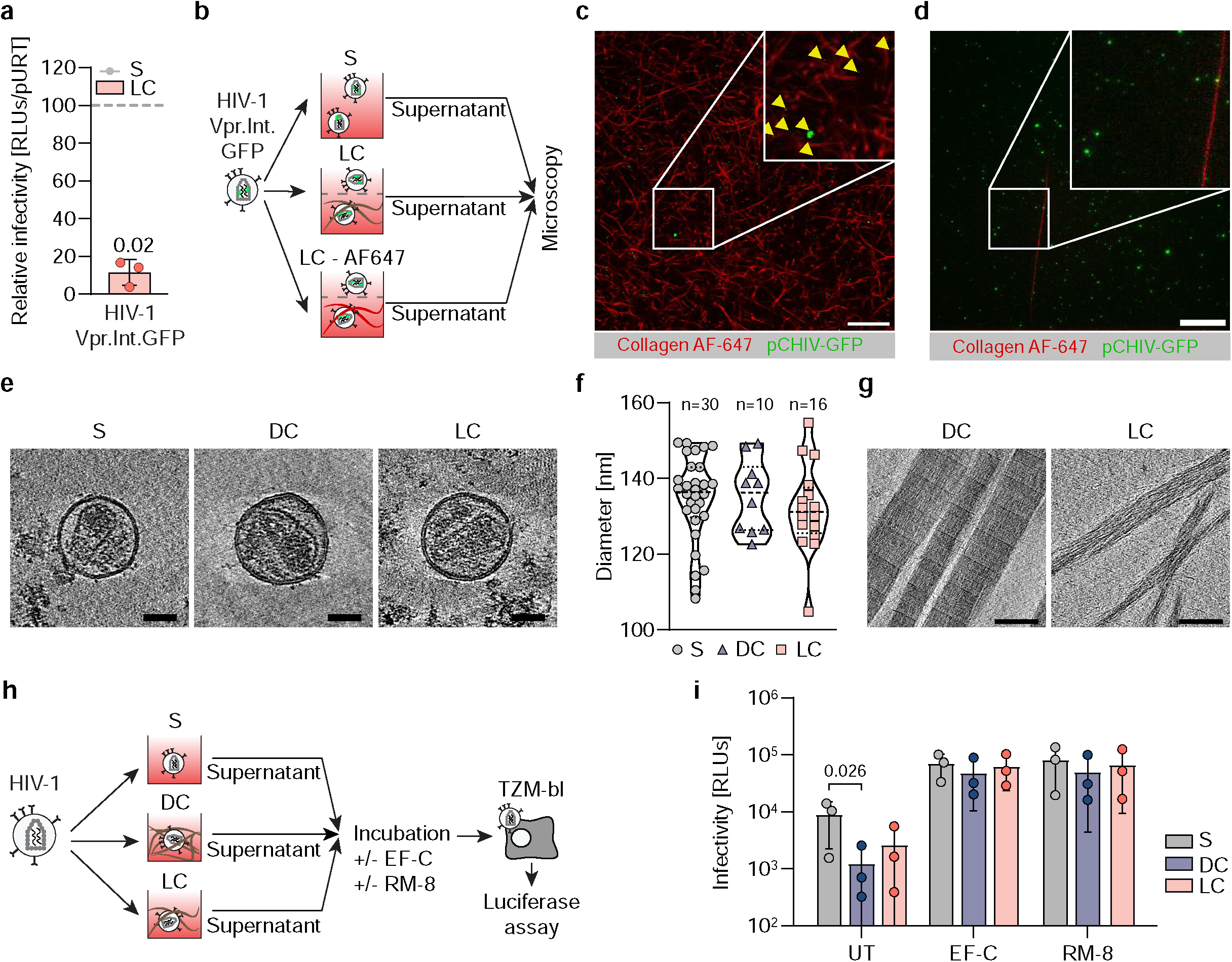
ERVI does not result from collagen deposition or structural damages of embedded virions. **a**. Relative infectivity of HIV-1 NL4.3 virus containing Vpr.Int.GFP fusion proteins after exposure to LC. **b**. Experimental workflow to assess collagen deposition at the surface of virions. HIV-1 NL4.3 Vpr.Int.GFP virions were collected from S, LC or LC-AF647 cultures and analyzed by microscopy. **c**. Representative confocal micrographs of HIV-1 NL4.3 Vpr.Int.GFP virions within AlexaFluor 647 stained LC matrices. Yellow arrows depict fluorescent virions. Scale bar: 20 µm. **d**. Representative TIRF-micrographs of HIV-1 NL4.3 Vpr.Int.GFP virions harvested from the supernatants of matrices in a. Scale bar: 5 µm. **e**. Averaged computational slices of a tomogram showing HIV-1 NL4.3 virions after 16h of culture in S (left panel), DC (middle panel), or LC (right panel). Scale bar = 50 nm. **f** Quantification of the diameter of virus particles treated as indicated in d. Violin plot shows individual data points with corresponding median, 25% and 75% quartiles. **g** Averaged computational slices of a tomogram showing disrupted dense (left panel) or loose (right panel) collagen fibers. Scale bar = 100 nm. **h**. Experimental workflow. HIV-1 NL4.3 virions retrieved from suspension or collagen matrices after 16h of culture were harvested, and equivalent amounts of RT units were incubated with infectivity enhancing peptides prior to infection of TZM-bl reporter cells. **i**. Relative infectivity of PNF treated virions. Results represent the mean ± SD of 3 independent experiments. Symbols indicate individual experiments. Significance is indicated by p-values, and was calculated by unpaired t-tests (d), or by two-way ANOVA test with Geisser-Greenhouse correction; Tukey’s post-test (h).

### ERVI is manifest at the step of virus fusion without affecting virion binding to target cells

The finding that infection enhancers boost the infectivity of HIV-1 particles subjected to ERVI suggested that ERVI acts at the early steps of the viral life cycle. To test if ERVI impairs the ability of HIV-1 particles to bind to target cells, we generated virions that incorporated Vpr.mRuby2 during virus production for visualization^40^ (Fig. 3a). Incorporation of Vpr.mRuby2 did not affect their sensitivity to infectivity reduction upon contact with 3D collagen (Fig. 3b). To visualize their interaction with the surface of TZM-bl target cells by spinning disk microscopy, cells were incubated with virus particles for 2h at 4°C to avoid particle internalization and to detect individual fluorescent HIV-1 particles attached to the cell surface (Fig. 3c). While incubating these particles for 16 h in suspension slightly reduced the number of virions detected at the surface of target cells (Fig. 3d, 2 +/- 0.5 bound virus particles/cell for fresh virus vs. 1.1 +/- 0.4 bound virus particles/cell for suspension), no additional reduction in binding efficacy was observed for virions that had been embedded in dense or loose 3D collagen. Notably however, prior contact with DC fibers resulted in significantly larger aggregates of virus particles being visible at the surface of target cells (suspension: 1.05+/-0.03 µm², DC: 3.47+/-0.8 µm²) (Fig. 3e). Such particle aggregation was not observed upon contact with LC (2.0+/- 0.5 µm²), indicating that this effect is not essential for ERVI. The predominant action of ERVI therefore is not at the level of virus binding to target cells.

**Figure 3:**
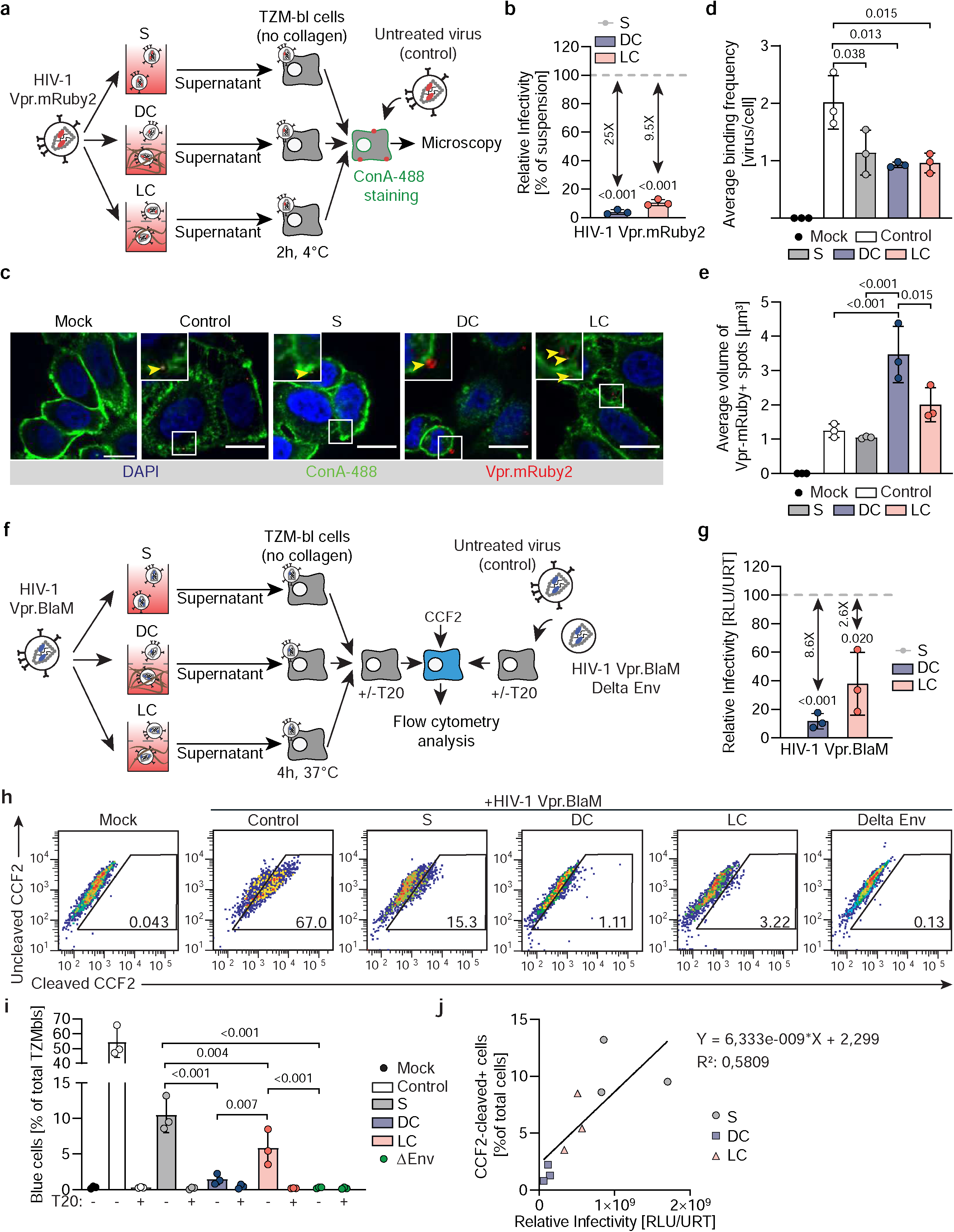
ERVI impairs HIV-1 entry but not binding to TZM-bl target cells. **a**. Experimental workflow. HIV-1 NL4.3 Vpr.mRuby2 virions were cultured in suspension or in collagen for 16h. Equivalent amounts of RT units from culture supernatants or untreated virions were then incubated with TZM-bl cells for 2h at 4°C. Cell membranes were then stained with Concanavalin-A AF-488, prior to microscopy processing. **b**. Relative infectivity of HIV-1 NL4.3 Vpr.mRuby2 virions after 16h of culture in suspension or collagen. **c**. Representative spinning disk micrographs of cells incubated with HIV-1 NL4.3 Vpr.mRuby2 virions. Yellow arrows indicate Vpr.mRuby2+ spots detected at the surface of the target cells. Scale bar: 15 µm. **d**. Average binding frequency of HIV-1 NL4.3 Vpr.mRuby2 virions to target cells as shown in c. **e**. Average volume of Vpr.mRuby2 spots detected as shown in c. **f**. Experimental workflow. HIV-1 NL4.3 Vpr.BlaM virions were cultured in suspension or in collagen for 16h. Equivalent amounts of RT units from culture supernatants or untreated virions were then incubated with TZM-bl cells for 4h at 37°C, in the presence or absence of T20 fusion inhibitor. Cells were then loaded with the CCF2-AM dye for 10h at 11°C and processed by flow cytometry. **g**. Relative infectivity of HIV-1 NL4.3 VprBlaM virions after 16h of culture in suspension or collagen. **h**. Representative flow cytometry dot plots of CCF2 loaded cells. Gates identify cleaved-CCF2+ cells. **i**. Quantification of the percentage of CCF2-product positive cells measured by flow cytometry. **j**. Correlation between Relative Infectivity and levels of CCF2-product positive cells after ERVI by linear regression. Results represent the mean ± SD of 3 independent experiments. Symbols indicate individual experiments. Significance is indicated by p-values, and was calculated by one-way ANOVA test; Dunnets’s post-test (b,g), or by one-way ANOVA test; Tukey’s post-test (d,e,i).

To assess the ability of HIV particles to fuse with target cells, Vpr.BlaM containing particles were produced and used to quantify the conversion of the ß-lactamase (BlaM) substrate CCF2 in the cytosol of target cells by flow cytometry as a measure for fusion ^40, 41^ (Fig. 3f). The incorporation of Vpr.Blam did not affect the sensitivity of the virions to ERVI in DC, while the infectivity reduction by LC was slightly less pronounced (Fig. 3g). Analyzing the fusion capacity of these particles revealed efficient cytosolic delivery of Vpr.Blam by HIV-1 particles kept in suspension (10.4 +/- 2.4%), which was dependent on the viral glycoprotein Env and could be inhibited by the HIV fusion inhibitor T20 (Fig. 3h,i). This fusion capacity was strongly impaired for particles that had prior contact with DC (1.5 +/- 0.7 %). LC (5.4 +/- 2.5%) had a less pronounced effect on the fusion capacity of HIV-1 particles and overall, infection rates and fusion capacity under the different conditions analyzed were correlated, albeit with deviations in particular for virions from suspension cultures (Fig. 3j). We conclude that the interaction of HIV-1 particles with tissue-like environments reduces their infectivity by impairing their ability to fuse with target cells and that ERVI may also affect additional post entry steps.

### ERVI moderately reduces infection of primary CD4+ T cells and MDMs

HeLa-derived TZM-bl cells are a convenient and widely used reporter cell to quantify the infectivity of HIV-1 particles but cannot reflect the differences in entry binding and receptor densities, membrane lipid composition as well as uptake pathways between different primary target cells^42^. We therefore analyzed the relevance of ERVI for infection of primary human CD4^+^ T cells and primary human monocyte-derived macrophages (MDMs) with a HIV-1 variant that uses CCR5 as entry coreceptor (HIV-1 NL4.3 R5) (Fig. 4a). This co-receptor tropism did not affect the sensitivity of these particles to ERVI when assessed on TZM-bl cells (Fig. 4b), which was again more pronounced in DC than LC. Productive infection of primary target cells was assessed by quantifying the percentage of cells with intracellular p24 capsid by flow cytometry (CD4^+^ T cells, Fig. 4c, MDMs, Fig.4d). The reverse transcriptase inhibitor efavirenz (EFZ) was used to define background detection of input virus. On both, CD4^+^ T cells and MDMs, productive infection by particles with prior contact to dense 3D collagen was significantly reduced (3.6-fold reduction to 27.5% of suspension on CD4^+^ T cells, 2.9-fold reduction to 34.8% of suspension on MDMs) and LC only mediated a very mild reduction that did not reach statistical significance (1.8-fold on CD4^+^ T cells, 1.4-fold on MDMs). These results reveal that ERVI reduces the infectivity of cell-free HIV-1 particles on primary target cells but with lower magnitude than on TZM-bl cells.

**Figure 4:**
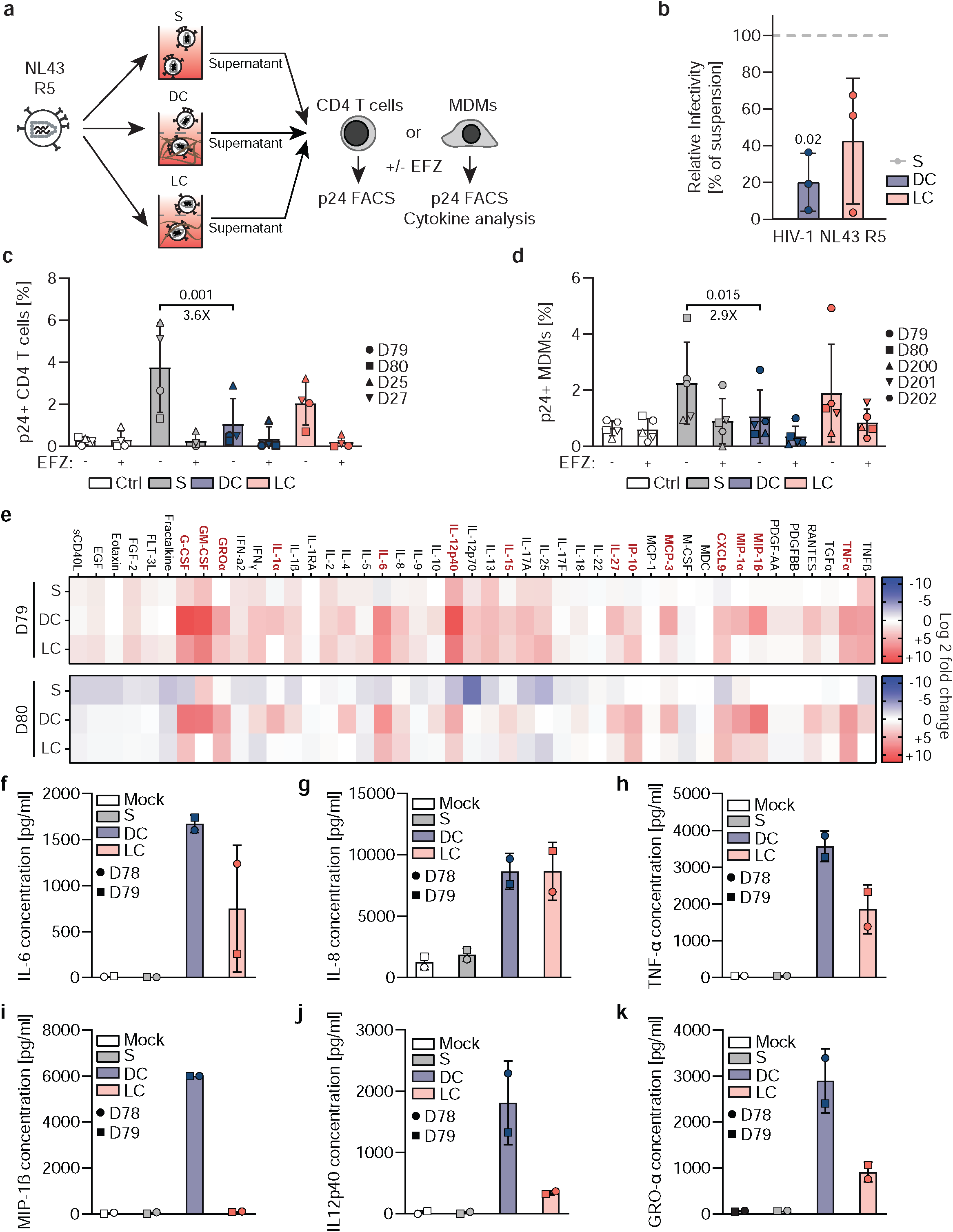
ERVI moderately restricts the infection of primary cells and sensitizes virus particles for innate immune recognition by MDMs. **a**. Experimental workflow. HIV-1 NL4.3 R5 virions were cultured in suspension or in collagen for 16h. Equivalent amounts of RT units were incubated with MDMs or activated CD4+ T cells, in the presence or absence of Efavirenz. Infection levels were determined by flow cytometry after p24 staining, and the supernatants from challenged MDMs were profiled for cytokine content. **b**. Relative infectivity of HIV-1 NL4.3 R5 virions after 16h of culture in suspension or collagen. Graphs depict mean values ± SD for n=3 experiments. **c**. Quantification of the percentage of p24+ CD4+ T cells by flow cytometry. Graphs depict mean values ± SD for 4 donors. **d**. Quantification of the percentage of p24+ MDMs by flow cytometry. Graphs depict mean values ± SD for 6 donors. **e**. Cytokine amounts in MDM supernatants 72h after challenge with virions from suspension or collagen cultures. Heatmaps indicate Log2 fold change of cytokine amounts in infected conditions as compared to mock infected conditions for 2 donors. Cytokines that were induced by DC cultured virions >5-fold as compared to suspension conditions for both donors are highlighted in red. **f-k.** Quantification of individual cytokines from the analysis shown in (e): IL-6 (**f**), IL-8 (**g**), TNF-α (**h**), MIP1β (**i**), IL12p40 (**j**) and GRO-α (**k**). Results represent the mean ± SD of 2-3 independent experiments. Symbols indicate individual donors. Significance is indicated by p-values, and was calculated by one-way ANOVA test; Tukey’s post-test (b,c,d).

With 27.5% infectivity remaining, the impact of ERVI on infection of primary CD4 T cells was less pronounced than on TZM-bl cells, where ERVI reduces virion infectivity to 14% of that of virions in suspension culture. Our computational model of HIV-1 replication in 3D collagen cultures of primary human mononuclear cells, which predicted that cell-associated virus transmission largely dominates over cell-free infection in 3D collagen matrices^7^, was based on the value of 14% of cell-free infectivity remaining in 3D. We therefore asked if the fact that the infectivity of these virions is higher on the primary target cells present in these 3D cultures affects this conclusion. To this end, we revisited our previous analyses estimating the efficacy of cell-free and cell-to-cell transmission within suspension and 3D collagen matrices by mathematical modelling^7^. Varying the parameter defining the reduced infectivity of cell-free infection within collagen compared to suspension, we estimate that the contribution of cell-free transmission to viral transmission within collagen continuously increases with increasing infectivity preservation (Sup Fig. 2a). Differences between estimates for LC and DC are partly affected by technical compensations within the fitting procedure due to model constraints, which lead to reduced estimates of cell-to-cell transmission rates within DC and thereby potentially underestimating the contribution of cell-to-cell transmission for this environment (Sup Fig. 2b). Under these conditions, the best estimates predict that an infectivity reduction to 27.5% results in a contribution of cell-cell transmission to overall virus spread of ∼60% and ∼30% in LC and DC, respectively. We conclude that in our primary cell 3D cultures, cell-associated transmission remains an important driver of HIV-1 spread in 3D collagen even if the infectivity of cell-free particles is less reduced than previously thought.

Since the effect of ERVI was much less pronounced upon infection of primary cells and MDMs in particular, we asked if ERVI has functional implications in MDMs in addition to reducing virion infectivity. Since MDMs can produce significant amounts of cytokines as consequence of innate immune recognition of HIV particles^43, 44, 45^, we quantified the amounts of a panel of cytokines in the supernatant of MDMs 3 days after challenge with HIV-1 particles with or without 3D collagen experience. Interestingly, prior encounter of HIV-1 particles with tissue-like 3D environments markedly and broadly altered the cytokine response of MDMs, resulting in increased release of important pro-inflammatory cytokines (Fig. 4e, compare suspension vs. DC and LC). Among the most differentially secreted cytokines were IL-6 (343-fold induction in DC, 154-fold in LC) (Fig. 4f), IL-8 (4.68-fold induction in DC and LC) (Fig. 4g), TNF-α (72-fold induction in DC, 38-fold in LC) (Fig. 4h), MIP-1b (82-fold induction in DC, 1.5-fold in LC) (Fig. 4i), IL12p40 (134-fold induction in DC, 2.26-fold in LC) (Fig. 4j) or GRO-α (39-fold induction in DC, 12-fold in LC) (Fig. 4k). Since no secretion of pro-inflammatory cytokines by MDMs was observed upon exposure of MDMs with culture supernatants of DC and LC matrices polymerized in the absence of HIV (Sup. Fig 3a) and endotoxins were undetectable in supernatants of 3D matrices polymerized in the absence of HIV (Sup. Fig. 3b), ERVI-induced sensing is not triggered by soluble matrix components or endotoxin contamination. ERVI thus sensitizes cell-free HIV-1 particles for specific innate immune recognition by MDMs.

### ERVI-induced innate immune recognition occurs during abortive infection and depends on the Env glycoprotein and the viral genome

Multiple HIV-1 components can be recognized by different PRRs in innate immune cells in the post entry phase of the viral life cycle (Fig. 5a). To identify the viral components that are recognized by cellular sensors after the collagen experience of HIV-particles, we challenged MDMs with virions derived from suspension or 3D cultures in the presence of selective inhibitors. These analyses focused on HIV-1 particles derived from DC matrices as they triggered the most pronounced sensing of HIV-1 particles. Interestingly, interfering with virus fusion by the entry inhibitor T20 or with EFZ did not impair the ERVI-mediated increase in IL-6 secretion (Fig. 5b). Entry and early post-entry steps of the HIV-1 life cycle are thus not required for ERVI-mediated induction of pro-inflammatory cytokine production, suggesting that innate recognition occurs in the context of non-productive, abortive, uptake of virus particles. To test for a direct involvement of the viral glycoprotein Env, we challenged MDMs with HIV-1 NL4.3 R5, a non-infectious variant lacking Env (HIV-1 ΔEnv) virions, or HIV-1 ΔEnv pseudotyped with VSVg (see Sup Fig. 4a-c for infectivity and glycoprotein incorporation). In contrast to virions carrying HIV-1 Env, virions pseudotyped with VSVg or lacking Env did not result in the secretion of IL-6 by MDMs after a collagen experience (Fig. 5c). The Env glycoprotein is thus necessary for the ERVI induced immune sensitization of HIV-1 particles. To test if the HIV-1 gRNA is required for innate sensing of collagen-experienced HIV-1 particles, MDMs were challenged with lentiviral particles with or without gRNAs and pseudotyped with HIV-1 Env or VSVg following exposure to collagen or medium. Only lentiviruses containing both HIV-1 gRNA and Env glycoproteins induced higher levels of IL-6 release after collagen imprint (Fig. 5d, see Sup Fig. S4d for IL-8 production). This increase in innate immune recognition of HIV-1 particles was not antagonized by increasing the local concentration of virion in the 3D environment (MOI 1: 2.7-fold induction for DC virus, MOI 10: 4.7-fold induction by DC) (Fig. 5e). Contrary to the effect of ERVI on virion infectivity, the capacity of the collagen matrix to sensitize HIV particles for innate immune recognition is therefore not saturable in the concentration range tested. Together, these results identify that the innate immune recognition of HIV-1 particles with prior contact with 3D collagen occurs in the course of abortive infections and involves the recognition of HIV-1 Env as well as HIV-1 gRNA.

**Figure 5:**
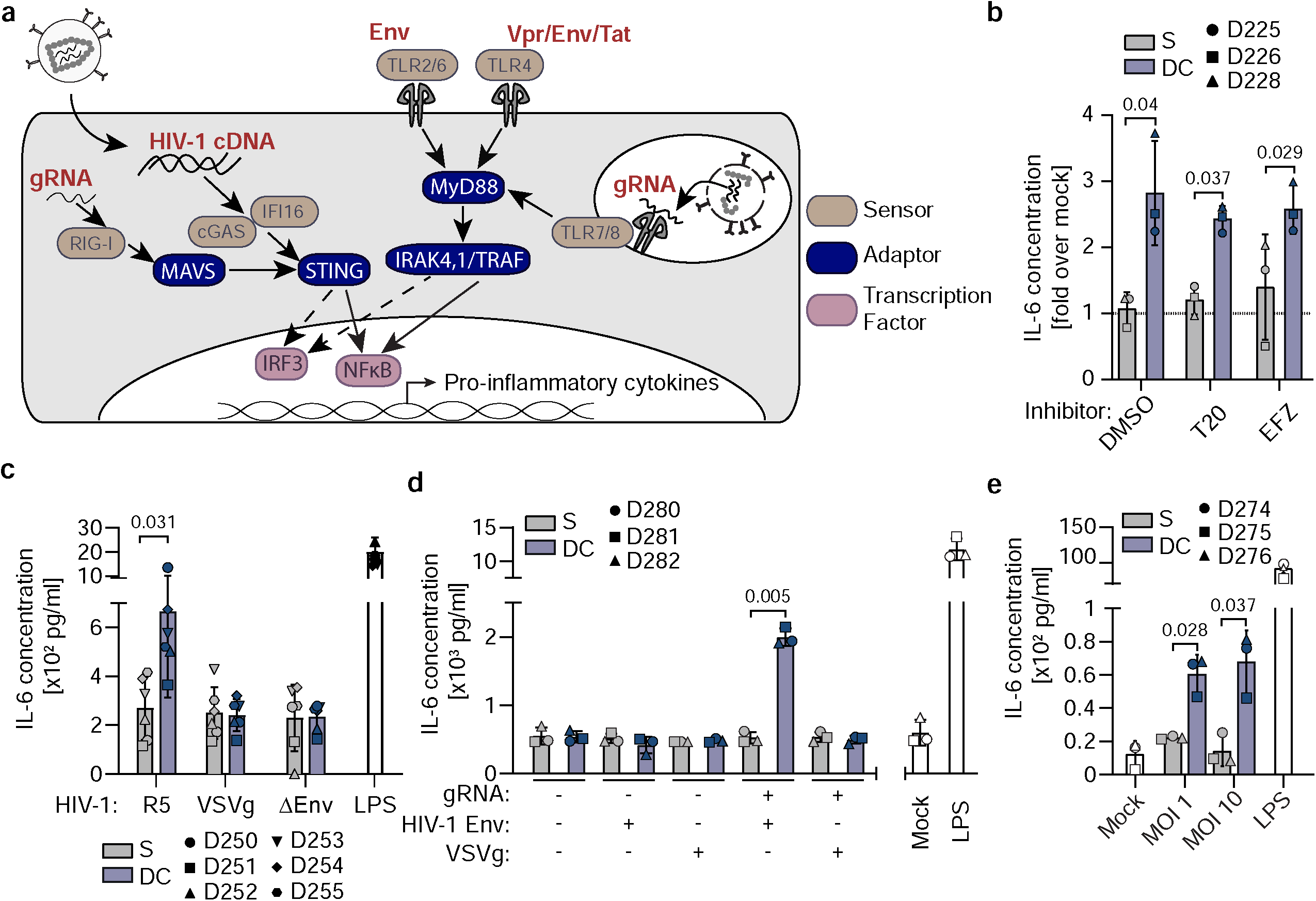
The collagen-induced innate immune sensitization of HIV-1 virions is dependent on the presence of Env and the viral genome. **a**. Schematic representation of innate immune sensors shown to detect HIV-1 derived PAMPs. PRRs: light brown, adaptor proteins: blue, PAMPs: red, transcription factors: purple. **b**. Measurement of IL-6 release by ELISA from the supernatants of MDMs after challenge with virions harvested from suspension or collagen cultures in the presence or absence of the T20 or EFZ HIV-1 inhibitors. Data is normalized to mock infected condition (grey dotted line). **c**. Measurement of IL-6 release by ELISA from the supernatants of MDMs challenged with NL4.3 R5, HIV-1 ΔEnv VSVg, or HIV-1 ΔEnv after culture in suspension or in collagen. **d**. Measurement of IL-6 release by ELISA from the supernatants of MDMs challenged with either naked, gRNA containing and/or VSVg/HIV-1 Env pseudotyped lentiviral particles cultured in suspension or in collagen. **e**. Measurement of IL-6 release by ELISA from the supernatants of MDMs after challenge with suspension or collagen experienced virions. Increasing amounts of virions were cultured in similar volumes in suspension or in collagen cultures, and equivalent amounts of virions harvested from these cultures were then used to challenge MDMs prior to ELISA analysis. Results represent the mean ± SD of 3 (b,c,e) or 6 (d) independent donors. Symbols indicate individual donors. Significance is indicated by p-values, and was calculated by paired t-tests or Wilcoxon matched-pairs signed rank test.

### Collagen-experienced HIV-1 virions are sensed by TLR 2/6 and routed into TLR-8 positive endosomes

To assess which PRRs are involved in the recognition of collagen-experienced HIV-1 particles (Fig. 6a), we tested the effect of PRR inhibitors on cytokine production of MDMs in response to challenge with HIV-1 particles subjected to ERVI. In line with the finding that fusion and post entry steps are dispensable for sensing (Fig. 5b), inhibition of the cytoplasmic DNA sensor cGAS had no effect on the ERVI-mediated induction of IL-6 or IL-8 secretion (Fig. 6b). Similarly, TLRs 1/2, 3 and 9 were dispensable for the recognition of ERVI-treated HIV particles. In contrast, inhibiting the general TLR signaling adaptor MyD88^46^, the endosomal TLR-8 that recognizes single stranded RNA^47^, or the use of GIT27, which impairs the activity of TLR-4 and TLR-2/6 fully abrogated the induction of IL-6 production by ERVI-treated HIV-1 (Fig. 6b, Sup Fig. 5a). Analyzing the full panel of secreted cytokines revealed that inhibition of either TLR-8 or TLR-4/TLR2/6 was sufficient to suppress the broad induction of proinflammatory cytokine production with no significant additional inhibition upon simultaneous use of both inhibitors (Sup Figs.5b-e). Using more selective inhibitors allowed to distinguish between TLR-2/6 and TLR-4 and identified a specific requirement for TLR-2, which recognizes diacylated lipopeptides^48, 49^ but also viral proteins^50, 51^ while TLR-4 was dispensable for the detection of collagen-experienced HIV-1 particles (Fig. 6c). These results suggest that (i) the increased production of pro-inflammatory cytokines by MDMs challenged with HIV-1 particles sensitized by 3D collagen reflects the recognition of HIV-1 Env by TLR-2/6 and gRNA by TLR-8 and (ii) both sensing events occur in one linear pathway.

**Figure 6:**
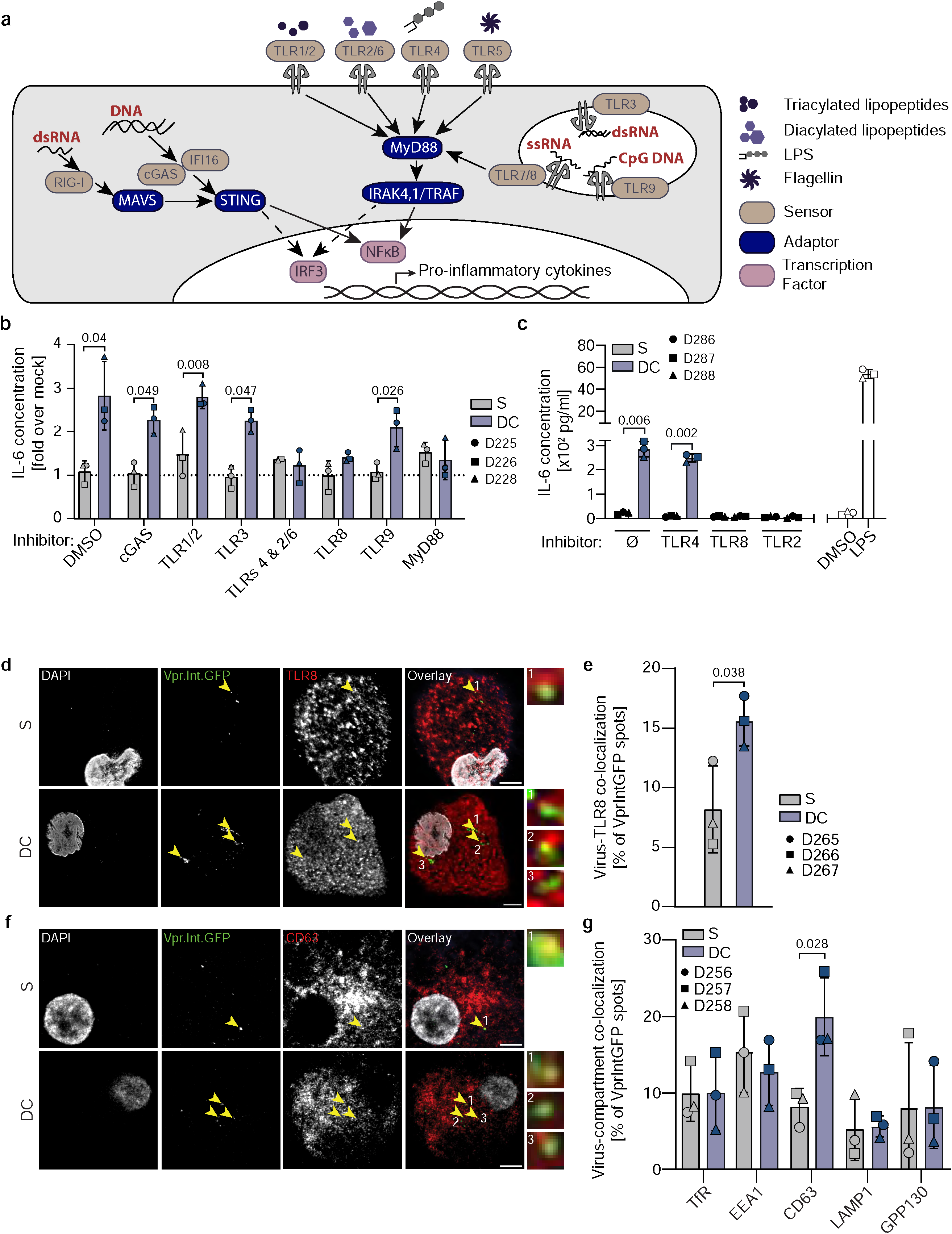
Collagen experienced HIV-1 virions are sensitized for TLR-2 and TLR-8 recognition in MDMs. **a**. Schematic representation of PRRs and their cognate ligands. PRRs: light brown, adaptor proteins: blue, PAMPs: red, transcription factors: purple. **b**. PRR inhibitor screening. MDMs were challenged with virions harvested from suspension or DC cultures in presence or absence of PRR inhibitors. IL-6 release was measured by ELISA from MDM supernatants. Data is normalized to mock infected condition (grey dotted line). **c.** Measurement of IL-6 release from the supernatants of MDMs challenged with HIV-1 virions retrieved from suspension or collagen cultures in presence or absence of the indicated inhibitors. **d**. Representative micrographs. MDMs challenged with HIV-1 NL4.3 R5 Vpr.Int.GFP virions after culture in suspension or in DC. Yellow arrows indicate Vpr.Int.GFP/TLR8 colocalization. Scale bar: 5 µm. **e**. Quantification of Virus/TLR8 colocalization from micrographs as in d. **f**. Representative micrographs. MDMs were challenged as in **d**. and immunostained for different compartments (CD63 shown). Yellow arrows: Vpr.Int.GFP/CD63+ colocalization. Scale bar: 5 µm. **g**. Quantification of Virus-compartment colocalization from micrographs as in f. Results represent the mean ± SD from 3 independent donors. Significance is indicated by p-values, and was calculated by paired t-tests or Wilcoxon matched-pairs signed rank test.

To gain more insight into this process, we followed the interaction of fluorescent HIV-1 Vpr.Int.GFP particles with MDMs and tested if the aggregation observed for HIV-1 particles derived from DC drives proinflammatory cytokine production. However, the induction of particle aggregation at the cell surface by incubation with RM-8 or EF-C did not increase innate immune sensing nor affect the sensitization of DC derived particles for sensing (Sup Figs. 6a-e). In contrast, the colocalization of HIV-1 particles following uptake by MDMs revealed that DC-derived HIV particles displayed significantly higher colocalization with intracellular vesicles positive for TLR-8 (Figs. 6d,e) or the endosome marker CD63 (Figs. 6f,g). Interactions with ECM thus primes HIV-1 particles for uptake into TLR-8 positive endosomes of MDMs, where the impact of ERVI may also affect the efficacy of gRNA sensing.

### A conformational change in Env is associated with increased innate immune recognition of DC-derived HIV-1 particles

The above results suggested that physical contact of HIV-1 particles with collagen fibers triggers the recognition of virions by TLR-2. HIV-1 Env adopts a complex trimeric structure that persist in a pre-triggered non-fusogenic conformation protected from neutralizing antibody recognition^52^. Subsequent binding to the primary receptor CD4 and co-receptor trigger rapid and pronounced structural rearrangements to facilitate fusion of the viral envelope with target cell membranes^53, 54^. Binding sites for antibodies and receptors on HIV-1 are well known (Fig. 7a) and the recognition of conformation-sensitive epitopes can be used to approximate the conformational state of Env. We therefore incubated suspension or DC-derived HIV-1 NL4.3 R5 particles with incorporated Vpr.Int.GFP with such antibodies or soluble CD4 (sCD4), detected these with a secondary red fluorescent reagent, and analyzed them by single particle microscopy (Fig. 7b). Accessibility of epitopes as indicated by colocalization of both fluorescent signals was significantly increased for DC-derived virions for sCD4 and the 17b and PG16 antibodies (Fig. 7c, d), indicating conformation changes at multiple surfaces of the Env trimer. LC-derived virions, which are not efficiently sensitized for innate recognition, failed to display similar conformation changes in Env (Sup Figs. 7a,b). We next compared the sensitization for innate immune recognition of Env in HIV-1 variants with different sensitivity to the ERVI infectivity reduction ranging from high (ADA), intermediate (CH167) to low (CH058.c). HIV-1 ADA was resistant to sensitization for innate immune sensing and CH058.c triggered the production of intermediate IL-6 levels. In stark contrast, DC-derived CH167 induced markedly more pronounced cytokine responses than HIV-1 NL4.3 (Fig. 7e, Sup Fig. 7c for IL-8 secretion). This increased susceptibility was associated with efficient routing of HIV-1 CH167 particles to TLR-8 positive endosomes (Figs. 7f, g, Sup. Movies 1: S, 2: DC). Collectively, our data suggest that interactions of HIV particles with ECM induce complex conformational changes in Env that affect its fusogenicity and recognition by TLR-2. The induction of TLR-2 recognition drives the internalization of particles into TLR-8 positive endosomes for sensing of gRNA that drives production of proinflammatory cytokines.

**Figure 7:**
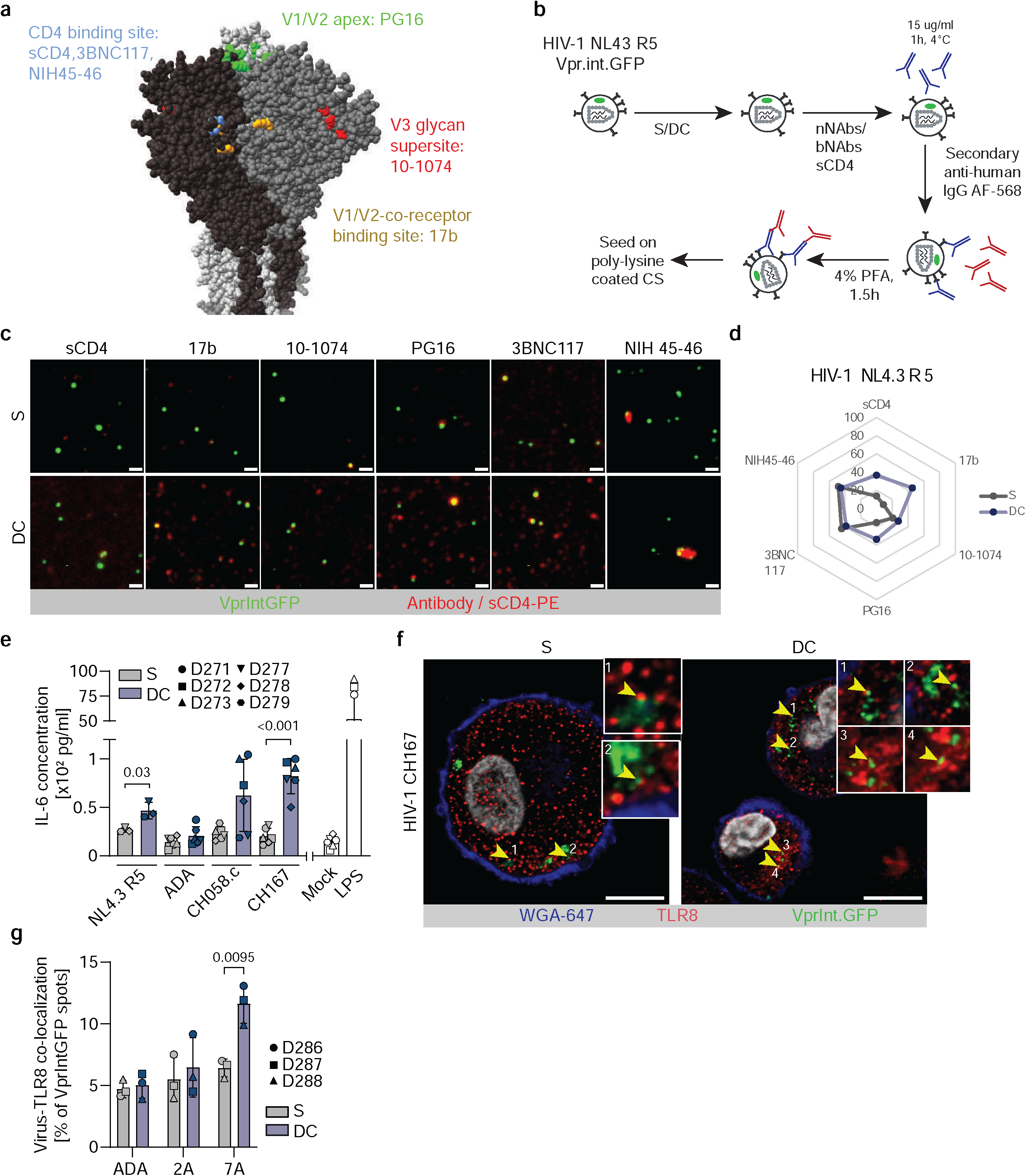
HIV-1 Env glycoproteins display differential accessibility for antibody binding after collagen contact. **a.** Representative structure of an HIV-1 Env trimer. Epitopes targeted by different bNAbs or nNAbs as well as sCD4 are highlighted (based upon a prediction of NL4.3 Env trimer structure using AlphaFold 3). **b**. Experimental workflow. HIV-1 NL4.3 R5 Vpr.Int.GFP virions harvested from suspension or collagen cultures were sequentially incubated with primary, epitope specific anti-gp120 antibodies, then with AF568 coupled secondary antibodies prior to processing for microscopy analysis. **c**. Representative micrographs of stained virions. Vpr.Int.GFP positive spots (green) can be seen bound by different antibodies or sCD4-PE (red). Scale bar: 0.5 µm. **d**. Spider plot representing the binding frequency of antibody binding to HIV-1 NL4.3 R5 as in (c). **e**. Measurement of IL-6 release by ELISA from the supernatants of MDMs challenged with different HIV-1 strains subjected to suspension or collagen cultures. **f**. Representative micrographs of MDMs challenged with HIV-1 CH167 virions cultured in suspension or in DC. Yellow arrows indicate Vpr.Int.GFP/TLR-8 colocalization. Scale bar: 10 µm. **g**. Quantification of virion/TLR-8 colocalization as in f. Results represent the mean ± SD of 3 independent experiments. Significance is indicated by p-values, and was calculated by paired t-tests.

## Discussion

The initial goal of this study was to define how tissue-like environments suppress the infectivity of cell-free virus particles. Investigating the mechanism and relevance of this intrinsic antiviral property of extracellular environments, ERVI, defined that this restriction affects the function of a broad range of viral glycoproteins without affecting glycoprotein incorporation and virion morphology. Since the infectivity restriction can be overcome by using infectivity enhancing PNFs, HIV-1 particles subjected to ERVI are not defective but display reduced infectivity. Mapping the effect in the viral life cycle defined virion fusion but not binding of cell-free virus particles with target cells as the key step affected by ERVI. Exposure to 3D collagen induced conformational changes in Env and the extent of these changes was reflected in the magnitude of the infectivity reduction. The transient contacts with collagen fibers thus alter the architecture of Env trimers in virions to reduce their fusogenicity.

Surprisingly, assessing the impact of ERVI on MDM target cells revealed a functional consequence of this extracellular restriction in addition to reducing virion infectivity: prior contact with tissue-like environments triggered innate immune recognition of HIV particles, resulting in the production of proinflammatory cytokines. Infection rates of ERVI-treated HIV particles do not correlate with cytokine production, blocking infection does not abrogate sensing, VSVg pseudotypes are reduced in their infectivity but not sensitized for innate immune recognition by ERVI, and infectivity reduction and sensitization for innate recognition differently react to alterations in matrix-particle ratios. Infectivity reduction and sensitization for innate immune recognition are thus mechanistically distinct consequences of ERVI. Our results demonstrate that this innate immune response reflects the sensing of Env by TLR-2 and the viral genome by TLR-8 with both sensing events being necessary for cytokine induction in the context of abortive, non-productive infections (Sup. Fig 8). The identification of a virus variant that is insensitive to the induction of innate recognition by 3D collagen allowed us to establish the following mechanistic model: in the absence of ERVI, only a minority of particles lead to productive infection while the majority of particles are subject to non-productive endocytic uptake^55, 56, 57, 58^. Some of these particles are sorted in TLR-8 positive endosomes but their gRNA is not efficiently sensed and particles are not detected by TLRs at the cell surface (Sup Fig. 8, left panel). Upon encounter of a tissue-like 3D environment, the conformational changes of Env reduce the infectivity of HIV-1 particles but also promote recognition of the viral glycoprotein by TLR-2, which increases sorting of particles into TLR-8 positive endosomes. In dendritic cells, HIV-1 hijacks the dynein motor protein SNAPIN to evade recognition by TLR-8^59^ and it will be interesting to dissect if TLR-2 signaling exerts similar effects in MDMs, possibly involving interactions of TLR-2 with SNAPIN^60^ However, the magnitude of the observed increase in colocalization of virions with TLR-8 does not seem to be sufficient to explain the substantial and broad induction of cytokine production. We therefore propose that recognition of HIV-1 particles by TLR-2 also has a qualitative impact on the efficiency by which TLR-8 can access viral gRNA in endosomes. Such an effect of TLR-2 signaling could affect the nature of the endosome, e.g. by regulation of its pH, to facilitate the access of TLR-8 to gRNA. In addition, the combined signaling capacities of TLR-2 and TLR-8 may be required in this scenario to trigger sufficient activation of MyD88 for marked downstream signaling. TLR8-mediated sensing of HIV-1 gRNA is associated with improved control of HIV replication in patients, the regulation of latency, and can be exploited by HIV to allow the productive infection of plasmacytoid dendritic cells^61, 62, 63, 64^. Similarly, TLR-2 has been implicated in the efficacy of HIV-1 spread, latency and chronic inflammation^65^. ERVI-mediated sensitization for innate recognition by TLR-2 and TLR-8 likely explains the role of these PRRs in HIV infection.

Our results reveal that the short and transient physical contact of HIV particles has significant impact on the fusogenicity of the viral glycoprotein Env as well as its interactions with TLR-2. The basal characterization of accessibility of epitopes in Env for antibody recognition suggests that ERVI drives conformational changes in Env with functional consequences. Since our results suggest conformational changes in Env as a key mechanism of ERVI, defining the structure-function relationship of this process will be an important goal of future studies. This will require high resolution structural analyses to define how ERVI-induced conformational changes in Env drive functional consequences in recognition by TLR-2 and infectivity reduction. The comparison of Env with differential sensitivity for the two ERVI effector functions will be an asset for such analyses. Interestingly, ADA Env was entirely resistant for sensitization to innate immune recognition and several Env proteins were only mildly impaired in their ability to mediate infection. Env therefore can, in principle, evade the antiviral activities of ERVI. Considering the multiple and complex roles of Env in HIV replication and pathogenesis, such adaptation likely comes with significant fitness cost and defining the balance between the two antiviral effects and virus spread in vivo warrants further investigation. Increased innate immune recognition of HIV-1 particles is also associated with the reduction of virion infectivity by the host cell factor SERINC5^44^. This restriction involves alterations in Env conformation^66, 67^ and both effects can be evaded by certain Env variants^68, 69, 70^. Regulation by the extracellular and intracellular adaptation of the viral glycoprotein thus emerges as an important parameter for innate immune responses to HIV-1 infection.

Our previous computational analysis had identified cell-associated infection as the predominant HIV-1 transmission mode in 3D cultures^7^. This conclusion was based on virion infectivity quantifications on model cell lines and was now questioned by our experimental finding that the effect of ERVI on infection of primary human CD4 T cells is less pronounced. Varying the magnitude of infectivity impairment in the model however revealed that even at these lower levels of cell-free infectivity reduction, cell-associated infection remains an important transmission mode. This relevant role of cell-associated transmission likely reflects that the long diffusion times required for virions to reach new target cells in 3D matrices are associated with significant reduction in their infectivity. This in turn raises the question why the production of large amounts of cell free virus particles is maintained in lentiviral evolution. The sensitization of non-infectious particles for the induction of pro-inflammatory cytokine responses by ECM identified in this study may be one reason since cytokines such as TNF-α, IL-2, IL-1 and IL-6 can increase the permissiveness of primary target cells to HIV-1 infection^71, 72, 73, 74^. Although the infection rates of local target cells are reduced by ERVI, the induced innate recognition may create a replication-prone tissue microenvironment and thereby indirectly facilitate virus spread in tissue. ERVI is exerted by 3D collagen from different species that form fibers of distinct architectures, suggesting that this restriction can be relevant in different HIV target tissues. Notably, chronic immune activation and inflammation observed in HIV patients even under therapy is associated with lymph node fibrosis, which contributes to HIV pathogenesis, e.g. by facilitating depletion of naïve CD4+ T cells^3, 75^. Fibrosis reflects the deposition of large amounts of collagen with altered structural and biophysical properties^76^, suggesting that the antiviral activity of ERVI is enhanced in fibrotic tissue. Since pro-inflammatory cytokines promote collagen deposition and thus fibrosis^77^, ERVI may significantly fuel fibrosis and CD4 T cell depletion.

Collectively, our study demonstrates that the biophysical properties of adhesive tissue-like environments exert a previously unrecognized, two-pronged antiviral mechanism that reduces virus spread by limiting virion infectivity and sensitizing virions for innate immune recognition. We propose that these tissue-intrinsic extracellular mechanisms synergize with the established cell-intrinsic mechanism to create antiviral innate immunity. In addition to its established role in supporting efficient adaptive immune responses in lymph nodes^78^, the ECM thus also emerges as an important regulator of antiviral innate immune responses.

## Materials and Methods

### Cells

293T cells (ATCC, CRL-3216) and TZM-bl reporter cells (courtesy of NIH AIDS Reagent Program (ARP-8129)) were cultured in Dulbecco’s Modified Eagle’s Medium (DMEM, Gibco) supplemented with 10% heat-inactivated fetal calf serum (FCS, Capricorn) and 1% penicillin/streptomycin (Gibco). Huh 7.5 cells were maintained in complete DMEM medium supplemented with non-essential amino acids.

### Primary CD4+ T cells and MDMs

Human peripheral blood of healthy, HIV-negative donors was obtained from the blood bank HD, according to regulation by local ethics committee (S-024/2022). CD4 T cells were isolated from human peripheral blood of healthy, HIV-negative donors using the RosetteSep Human CD4 T cell enrichment kit (15062, StemCell Technologies) according to the manufacturer’s protocol. The cells were then activated with Dynabead Human T-Activator CD3/CD28 (11131D, Gibco), or in a 3x3 activation as described previously^7^ for 72h and cultured in Roswell Park Memorial Institute Medium (RPMI, Gibco) supplemented with 10% heat-inactivated FCS, 1% penicillin–streptomycin, and 10 ng/ml interleukin 2 (IL-2, Biomol). For Monocyte-derived macrophages (MDMs), peripheral blood mononuclear cells (PBMCs) were isolated from buffy coats by Biocoll (Merck Biochrom) density gradient centrifugation. CD14^+^ monocytes were then isolated from PBMCs by positive selection using magnetic beads (CD14 MicroBeads; 130-050-201, Miltenyi Biotech) and an AutoMACS Pro Separator (Miltenyi Biotech). 1x10^5^ monocytes per well were then seeded in glass-bottom 96 well plates that were previously coated with fibronectin (2 µg/cm²) and maintained at 37°C with 5% CO_2_ in complete RPMI in the presence of 5% human AB serum (H4522, Sigma-Aldrich) for differentiation into MDMs for 10-14 day^44^.

### Viruses

Virus stocks of replication competent HIV-1 and HIV-2 strains (pNL4.3 WT, pNL4.3 ΔNef, pNL4.3 R5, ADA, CH077, CH0 58.c, CH0198, RHGA, CH0167, CH0293, HIV-2 Rod9-GFP) were generated by transfecting 293T cells (sub-confluent 15cm² dishes) with 25 µg of proviral constructs alongside 75 µl of linear polyethyleneimine (PEI, Sigma Aldrich) in Optimem medium (Gibco). Viruses containing Vpr-BlaM, Vpr.mRuby2 or Vpr.Int.GFP fusion proteins were similarly produced by PEI co-transfection of 293T cells using 25 µg of pNL4.3 provirus with 7.5 µg of Vpr fusion constructs. For single round HIV-1 virus production (pNL4.3 ΔEnv VSVg, pNL4.3 ΔEnv HIV-1 Env), 22µg of pNL4.3 ΔEnv were used, complemented with 3 µg of the respective glycoprotein expression vectors^40, 44^. Two to three days post transfection, supernatants were harvested, filtered (0.45 µm) and ultracentrifuged through a 20% (w/v) sucrose cushion. Virus pellets were then resuspended in sterile filtered PBS 0.1% BSA, aliquoted and stored at -80°C. All replication competent virions were handled in a BSL-3 containment laboratory.

### Lentiviral pseudotyping

Lentiviral stocks were prepared by PEI transfection of sub-confluent 293T cultures. Briefly, 22.5 µg of pWXPL-GFP lentiviral backbone was co-transfected alongside 2.3 µg of pAdvantage, 15 µg packaging vector Pax2 and 8 µg of envelope glycoprotein encoding plasmids (VSVg, pcDNA Con1, pcDNA JFH1). Lentiviral vectors carrying Vpx_mac239_ were produced in 293T cells by co-transfection of pWPI, pcDNA.Vpxmac239, pΔR8.9 NSDP, and VSV-G at a molar ratio of 4:1:3:1 ^44^. Cell culture supernatants were harvested and filtered 3 days post transfection, and further concentrated through a 20% (w/v) sucrose cushion ultracentrifugation. Aliquots were stored at -80°C. Virus titers were determined by SG-PERT analysis.

### Plasmids

The proviral plasmids pNL4.3 WT and ΔNef have been previously described^79^. The pNL4.3-R5 WT & ΔNef, containing 7 point mutations in the NL4.3 Env were previously described^44, 80^. The pNL4.3 ΔEnv, and HIV-2 Rod9-GFP proviruses were previously described^81^, provirus encoding the CCR5 tropic ADA provirus were via the NIH AIDS reagent program (ARP-416)^82^. All Transmitted/Founder strains were obtained via the NIH AIDS reagent program (pCR-XL-TOPO CH077 (ARP-11742)^83^, pCR-XL-TOPO CH058.c (ARP-11856)^83^, pCR-XL-TOPO_HIV-1 M subtype C CH198^84^. And chronic strains (pBR322 HIV-1 M subtype B STCO^85^, pBR322 HIV-1 M subtype B RHGA(ARP-12421)^85^, pUC57_HIV-1 M subtype C CH167(ARP-13544), pUC57rev_HIV-1 M subtype C CH293(ARP-13539)^84^. The following expression plasmids were used: pMM311 (Vpr.BlaM)^41^, the pmRuby2.Vpr vector was previously described^86^ and kindly provided by Dr. Tom Hope. The Vpr.Int.GFP expression plasmid was kindly provided by Anna Cereseto. The pcDNA3.1 Vpx SIVmac239-Myc and the pΔR8.9 packaging vector harboring a Vpx binding motif were previously described^81^.

### Reagents & antibodies

The following dyes were used: Concanavalin A-488 (C7642, Sigma Aldrich) was used a concentration of 50 µ/ml to stain TZM-bl cell membrane for 10 min at RT in the dark, Alexa Fluor 488 NHS Ester (A20000, Invitrogen) was used to fluorescently label loose collagen gels as described below, Fluoromount DAPI (00-4959-52, Invitrogen) was used to mount coverslips and stain cell nuclei, the CCF2-AM dye (K1023, Thermo Scientific) was used to perform HIV-1 entry assays, and WGA-647 (W32466, Invitrogen) was used according to manufacturer protocol to label MDM plasma membranes. The following reagents were used: PEI (Sigma Aldrich), Pur-A-Lyzer™ Mega 3500 kit (PURG35010, Sigma Aldrich). The following antibodies were used: Zombie Violet dye (Biolegend), anti-p24 KC-57 antibody (Beckman Coulter), rabbit anti-HIV-1 p24 (Kindly provided by Prof. Dr. Barbara Müller), rabbit anti-HIV-1 gp120 (Valerie Bosch, DKFZ, Heidelberg), mouse anti-TLR-8 (67317, Proteintech), mouse anti-EEA1 (68065, Proteintech), mouse anti-CD63 (556019, BD), sheep anti-TGN46 (AHP500GT, BioRad), mouse anti-GPP130 (923801, Biolegend), mouse anti-LAMP1 (ab25630, Abcam), mouse anti-TfR (13-6800, Invitrogen), anti-HIV-1 gp120 (PG16: ARP-12150; 3BNC117: ARP-12474; 10-1074: ARP-12477; NIH45-46: ARP-12174; 17b: ARP-4091) and sCD4-PE (CD4-HP2H8, ACRO Biosystems). Secondary antibodies used were, anti-mouse, anti-sheep or anti-human IgG1 AlexaFluor-568 antibodies (Invitrogen), as well as goat anti-rabbit IRDye700/800 conjugated antibodies (926-32211, Rockland), and goat anti-rabbit IgG-HRP (31460, Invitrogen).

### Virus titer determination

One step Syber Green based Product Enhanced Reverse Transcriptase assay (SG-PERT) was used to assess HIV-1 virus titers as described previously (Vermeire et al 2012). Briefly, concentrated virus stocks were first diluted in PBS, or culture supernatants were directly lysed in 2x lysis buffer (50 mM KCL, 100 mM Tris-HCl pH 7.4, 40% glycerol, 0.25% Triton X-100) supplemented with 40 mU/µl Rnase inhibitor for 10 minutes at room temperature. Lysed samples were then exported from BSL-3 and further diluted 1:10 in dilution buffer (5 mM (NH_4_)_2_SO_4,_ 20 mM KCL, 20 mM Tris-HCl pH 8). In parallel, 10 µl per well of a dilution series of a virus standard (pCHIV, 8,09x10^8^ pURT/µl) were also lysed for 10 min. All lysed samples were then incubated with 10 µl of 2x reaction buffer (1xdilution buffer, 10 mM MgCl_2_. 2x BSA, 400 µM each dATP, dTTP, dCTP, dGTP, 1pmol of each RT forward and reverse primers, 8 ng MS2 RNA, SYBR Green 1:10000) supplemented with 0.5U of GoTaq HotStart Polymerase. RT-PCR reactions were carried out and read in a real-time PCR detector (CFX 96, Biorad) using the following program: (1) 42 °C for .20 min, (2) 95°C for 2 min, (3) 95°C for 5s, (4) 60°C for 5 s, (5) 72°C for 15 s, 80°C for 7 s, repeat steps 3 to 6 for 40 cycles with a melting curve read out as a final step. Sequence of the used primers are as follows: fwd RT primer (TCCTGCTCAACTTCCTGTCGAG) and rev RT primer (CACAGGTCAAACCTCCTAGGAATG.)

### Virus infectivity determination

To assess the infectivity of virus stocks, TZM-bl reporter cells were infected as previously described ^87^. In brief, TZM-bl cells stably expressing HIV-1 entry receptors and containing luciferase and β-galactosidase genes under the control of the HIV-1 LTR promoter were infected using dilution series of concentrated virus. 72h post-infection, cells were fixed in 3% PFA, and incubated with a substrate solution (β-Gal supplemented with 200 µg/ml X-Gal) for 3h at 37°C. Blue cells were counted to determine infectious virus titers in the form of Blue Cell Units (BCUs). For virus containing cell culture supernatants, TZM-bl cells were infected in triplicates, and were lysed 3 days post-infection using 1x lysis buffer (Promega) for 10 min. Lysates were then incubated with Luciferase substrate (Promega), and luciferase activity was measured for 5s at a Tecan Infinity luminometer.

For saturation experiments, 10^5,^ 10^6^, or 10^7^ BCUs were used for seeding or embedding in collagen matrices of equivalent volumes. The culture supernatants were then processed by SG-PERT. TZM-bl cells were infected using equivalent amounts of RT units for all conditions. Virus infectivity after seeding or embedding at different concentrations was then determined by measuring the average luciferase activity from cell lysates 3 days post infection. To assess the relative infectivity of viruses after seeding or embedding, the infectivity and RT activity of virus containing supernatants were assessed as described above.

The relative infectivity of the respective supernatants was calculated as the average luciferase activity divided by the average amount of RT units measured from the culture supernatant.

### Virus embedding in 3D matrices

Type I collagen matrices of different densities were polymerized as previously described ^7^. In brief, dense collagen gels (3 mg/ml) were generated by mixing 10X MEM medium with 7.5% NaHCO_3_ (both Gibco) and highly concentrated rat tail collagen I (354236, Corning) at a 1:1:8 ratio on ice. Concentrated virus in complete DMEM was then added to the neutralized and chilled collagen solution at a 1:1 ratio. 100 µl of virus containing collagen solutions were then distributed in each well in a 96 well-plate and allowed to polymerize within 15 minutes at 37°C. Loose collagen gels (1.7 mg/ml) were generated by mixing 10X MEM medium with 7.5% NaHCO_3_ (both Gibco) and PureCol bovine skin collagen I (5005, Advanced Biomatrix) at a 1:1:8 ratio on ice. Concentrated virus was then added to the neutralized and chilled collagen solution at a 1:1 ratio and, and the mixture was allowed to pre-polymerize for 10 min at 37°C. 100 µl per well were then transferred to a 96-well plate. Gels were allowed to polymerize within 45 minutes at 37°C.

Human lyophilized placental type III collagen (5019, Advance Biomatrix) was reconstituted using a chilled 2 mM CH_3_COOH solution on ice to reach 6 mg/ml. Type III collagen matrices (3 mg/ml) were generated by mixing 10X MEM with 7.5% NaHCO_3_ (both Gibco) with the reconstituted type III collagen solution at a 1:1:8 ratio on ice. Concentrated virus in complete DMEM was then added to the neutralized and chilled collagen solution at a 1:1 ratio. Gels were allowed to polymerize within 30 min at 37°C.

Matrigel Growth Factor Reduced Basement Membrane (354230, Corning) gels (5 mg/ml) were generated according to manufacturer protocol. In brief, concentrated Matrigel aliquots (8.9 mg/ml) were thawed at 4°C overnight. Thawed Matrigel was then combined with complete DMEM containing concentrated virus at a 3:2 ratio on ice. 100 µl of mixture was transferred to a 96-well plate, gels were allowed to polymerize within 45 min at 37°C.

Agarose gels were prepared by preparing a 0.8% agarose solution in PBS. The dissolved agarose solution was combined 1:1 with complete DMEM containing concentrated viruses. 100 µl of the mixture was transferred to a 96-well plate, gels were allowed to polymerize at 4°C within 15 minutes.

All polymerized gels were then overlaid with 100 µl pre-warmed complete DMEM and incubated at 37°C. Culture supernatants were harvested at the indicated time points and processed for relative infectivity determination.

### Western blotting

For detection of gp120 on virions from supernatants of suspension and collagen cultures, supernatants were concentrated by ultracentrifugation through a 20% sucrose cushion (44000 rpm, 45 min) using an Optima LE-80k ultracentrifuge (Beckman Coulter), and lysed in 2x SDS sample buffer (10% glycerol, 6% SDS, 130 mM Tris Hcl pH 6.8, 10% β-Mercaptoethanol) and boiled for 5 min. Protein samples and were resolved with SDS-PAGE electrophoresis and blotted onto nitrocellulose membranes. Membranes were blocked in 4% milk/TBST for 1 h and were incubated with primary antibodies overnight. We used secondary antibodies conjugated to IRDye700/800 (1:20000, Rockland) for fluorescent detection with Licor (Odyssey).

### Fluorescent collagen gels

Fluorescently labelled LC gels were generated as described previously^88^. In brief, 5 mg of Alexa Fluor 647 NHS Ester dye (A20006, Invitrogen) was dissolved in 0.5 ml DMSO. The dissolved dye was then combined with PureCol bovine skin collagen (3 mg/ml) at a 1:100 ratio on ice and stirred overnight at 4°C. After labelling, the excess dye was removed by dialysis using a 3.5 kDa molecular weight cut off Pur-A-Lyzer™ Mega 3500 kit (PURG35010, Sigma Aldrich), placed in 1 L of acetic acid solution (0.02 N, pH 3.9) and stirred for one week at 4°C. The resulting collagen solution was kept at 4°C until further use. Fluorescently labelled collagen gels were polymerized as described above, by combining labelled and unlabeled collagen solutions at a 1:10 ratio.

### Confocal reflection microscopy

Collagen gels were imaged by confocal reflection microscopy as previously described ^7, 89^. Briefly, different collagen gels were generated in 15 well angiogenesis µ-slides (Ibidi) as described above. Point laser scanning confocal microscopy was then performed on a Leica SP8 microscope using an HC PL APO CS2 63x/1.4 N.A. oil immersion objective. Images were acquired using PMT detectors in reflection mode with a laser excitation at 567 nm, and a spectral detection window set between 550 nm and 570 nm wavelengths. Fluorescently stained collagen matrices were additionally imaged using a 488 nm laser.

### Widefield imaging

Viruses were embedded in stained or unstained LC matrices as described above. The culture supernatants were then harvested and ultracentrifuged through a 20% (w/v) sucrose cushion. Virus pellets were resuspended in 3% PFA for 90 min. The fixed virus particles were then seeded on 0.01% poly-L-lysine coated coverslips for 30 min and mounted on microscopy slides with Fluoromount DAPI (Invitrogen). Samples were then imaged using an epifluorescence microscope (Olympus IX81 S1F-3) under a 100X oil objective (PlanApo, N.a. 1.40). Quantification of double positive Vpr-mRuby2 and Alexa Fluor 647 signals was performed using the Spot Detector plugin of the Icy image analysis software.

### TIRF microscopy

To exclude the possibility that collagen fragments remain attached at the surface of virions after contact with collagen fibers, HIV-1 NL4.3 R5 Vpr.Int.GFP virions were embedded in fluorescently labeled collagen for 16h. The supernatants were then harvested and transferred in 0.01% poly-L-lysine coated µ-Slide 8 Well (80821, Ibidi). TIRF microscopy of the samples was performed on a Zeiss Axio Observer microscopy stand using an alpha PlanAPO 100x/ 1.46 N.A. oil objective and a Teledyne Prime-BSI sCMOS camera (6.5 µm x 6.µm pixel size). The sample was excited with a 561 nm or a 488 nm Visitron VS-laser control (Visitron Systems GmbH) and emission light was collected using Chroma TIRF Quad Line Beamsplitter with Quad Line Rejectionband 405/488/561/640 or Laser Beamsplitter zt 488 with 525/50 band-pass filter, respectively. The TIRF illumination was realized by a full 360-degree laser beam rotation at the back focal plane using 2D galvo scanners (Ring-TIRF). The evanescent field penetration depth was adjusted to 200nm. Laser intensities, shutter, and camera were controlled using the VisiView program (Visitron Systems GmbH). The presence of AlexaFluor 647 labeled collagen fibers was assessed at the surface of Vpr.Int.GFP spots using the Icy software.

### Harvesting of collagen fibers for cryo-ET analysis

To assess the morphology of single collagen fibers by cryo ET, DC or LC gels were prepared as previously in 1.5 ml Eppendorf tubes. Polymerized gels were disrupted by sonication on a Sonorex super RK 102 H sonicator for 10 sec on ice. Disrupted individual fibers were resuspended in PBS and further processed for cryo-ET.

### Plunge freezing

Collagen fibers and supernatant used for plunge freezing were collected as described above. Holey carbon grids (Cu 200 mesh, R2/1, Quantifoil®) were plasma-cleaned for 10 s in a Gatan Solarus 950 (Gatan). Samples were mixed with 10x concentrated 10 nm protein A gold (Aurion) prior plunge freezing. A total volume of 3 µl was used for plunge freezing into liquid ethane using an automatic plunge freezer EM GP2 (Leica). The ethane temperature was set to −183 °C and the chamber to 24 °C with 80% humidity. Grids were blotted from the back with Whatman™ Type 1 paper for 3 s. Grids were clipped into AutoGrids™ (Thermo Fisher Scientific).

### Cryo-electron tomography and tomogram reconstruction

Cryo-electron tomography was performed using a Krios cryo-TEM (Thermo Fisher Scientific) operated at 300 keV and equipped with a post-column BioQuantum Gatan Imaging energy filter (Gatan) and K3 direct electron detector (Gatan) with an energy slit set to 15 eV. As a first step, positions on the grid were mapped at 8,700× (pixel spacing of 10.64 Å) using a defocus of approximately -65 µm in SerialEM ^90^ ^t^o localize collagen fibers or HIV particles. Tilt series were acquired using a dose-symmetric tilting scheme ^91^ with a nominal tilt range of 60° to -60° with 3° increments with SerialEM. Tilt series were acquired at target focus −4 μm, with an electron dose per record of 3 e^-^/Å^2^ and a magnification of 33,000× (pixel spacing of 2.671 Å). Beam-induced sample motion and drift were corrected using MotionCor2^92^. Tilt series were aligned using AreTomo ^93^ and tomograms were reconstructed using R-weighted back projection algorithm with dose-weighting filter and SIRT-like filter 5 in IMOD^94^. Tomograms were used to measure the diameter of HIV particles in IMOD. For visualization, tilt series were aligned using protein A gold as fiducials in IMOD. Tomograms in Fig. 2 were reconstructed using R-weighted back projection algorithm with 3DCTF, dose-weighting filter and SIRT-like filter 10. In IMOD, 15 slices of the final tomogram were averaged and Fourier filtered. The diameter of HIV particles was measured from the outer leaflet of the viral membrane in IMOD. The average diameter of two measurements per particle was used in the graph.

### Infectivity enhancement experiments

HIV-1 virions were cultured in suspension or in collagen for 16h. After performing an SG-PERT analysis from the culture supernatants, equivalent amounts of reverse transcriptase units were then incubated with the specified infectivity enhancers for 20 min at 37°C prior infection of TZM-bl cells. Concentrations of infectivity enhancers used: EF-C (15 µg/ml) ^37^ or RM-8 (15 µg/ml) ^38^ The relative infectivity of the treated virions was assessed as previously described.

### Binding assay

NL4.3 Vpr mRuby2 virions were cultured in suspension or in collagen for 16h. Viral titers in the supernatants were quantified by SG-PERT. Equivalent amounts of RT units were then used to infect TZM-bl cells seeded on glass coverslips in 24-well plates 24h prior to infection. After 2h at 4°C to prevent virion internalization, the cells were then washed with PBS, and the plasma membrane was stained with Concanavalin A Alexa Fluor-488 (Invitrogen) according to manufacturer protocol. The samples were then fixed using 3% PFA for 90 min, and mounted on microscopy slides using Fluoromount-DAPI (Invitrogen). Spinning disk confocal microscopy was performed on a PerkinElmer UltraVIEW VoX microscope equipped with a Yokogawa CSU-X1 spinning disk head and a Nikon TiE microscope body. An Apo TIRF 60x/1.49 N.A. oil immersion objective and a Hamamatsu C9100-23B EM-CCD camera were used. Images were acquired using solid state lasers with excitation at 405nm, 488nm and 561nm with matching emission filters. Z-stacks were acquired with a z-spacing of 0.5 µM steps. The percentage of cells with bound virus was then analysed using the Imaris software (Oxford Instruments) in which Z-stacks were reconstructed in 3D. Cell membranes and virions were segmented as individual surfaces, and statistic values (surface volumes, relative distance of surfaces) were retrieved from the software.

### Vpr.BlaM entry Assay

TZM-bl reporter cells were infected with an MOI of 0.1 using HIV-1 NL4.3 viruses containing Vpr-BlaM after treatment in suspension or collagen for 16 hours. Concentrated virus that was not seeded in medium prior to infection was used as positive control, virus particles lacking Env were used as negative control. Fusion of HIV-1 particles was allowed to proceed for 4 h at 37°C. Cells were then washed twice in PBS and stained with 2mM CCF2-AM dye (Invitrogen) supplemented with 2.5 mM Probenecid in Fluorobrite DMEM 2% FCS for 6 h at 11°C to prevent particle fusion during the staining process according to manufacturer instructions. Where indicated, T20 was added to block fusion. Cells were then trypsinized fixed in 3% PFA in PBS at 4°C overnight and analyzed by FACS.

### Primary CD4+ T cell infection

Activated primary CD4+ T cells were infected in 96-well plated in triplicate with suspension and collagen culture supernatants containing NL4.3 R5 as previously described ^7^. Briefly, equivalent RT units were used between the different conditions, as quantified by SG-PERT. The cells were spin-infected at 2000 rpm for 90 min in presence or absence of 3 µg/ml reverse transcriptase inhibitor Efavirenz (EFZ), then cultured at 37°C for 3 days. The samples were then stained with a fixable Zombie Violet dye (Biolegend), fixed in 3% PFA for 90 min, then stained with an anti-p24 KC-57 FITC antibody (Beckman Coulter) in 0.1% Triton 100-X for 30 min at 4°C according to manufacturer protocol. Samples were then measured by flow cytometry.

### Primary MDM infection

Differentiated macrophages were spin transduced at 37°C in a pre-heated centrifuge with lentiviral vectors containing Vpx_mac239_ for 1h at 300 rpm. Within 16h post-transduction, cells were infected in triplicates with equivalent amounts of RT units as measured by SG-PERT from suspension and collagen culture supernatants in a 96-well plate format, in presence or absence of EFZ. MDMs were also treated with 50 ng/ml LPS as positive control. Infected MDM culture supernatants were harvested 3 days post-infection and further processed for cytokine analysis. 5 days post-infection, cells were harvested by trypsinization, stained with a fixable Zombie Violet dye (Biolegend), fixed in 3% PFA for 90 min, then stained with an anti-p24 KC-57 FITC antibody (Beckman Coulter) in 0.1% Triton 100-X for 30min at 4°C according to manufacturer protocol. Infection rates were then determined by flow cytometry.

### Cytokine quantification

The amounts of cytokines and chemokines present in cell culture supernatants were determined by Eve Technologies Corporation using the Discovery Assay®: Human Cytokine Array/Chemokine Array 48-Plex. Samples were exported out of BSL-3 containment after treatment with 0.5% Triton 100-X for 30 min prior to shipping. Results are in pg/ml of cytokines/chemokines according to the company protein standard. Cell-free supernatants were also analyzed for levels of Il-6, Il-8 and TNF-α by enzyme-linked immunosorbent assay (OptEIA ELISA kits; BD Biosciences) according to manufacturer’s instructions.

### Endotoxin level determination

To exclude potential endotoxin contamination of our collagen stocks, supernatants harvested from agarose, type I or III collagens, as well as matrigel matrices (polymerized in the absence of virus) were used for endotoxin testing. Endotoxin levels were quantified using a Pierce Chromogenic Endotoxin Quant Kit (ThermoScientific, A39553) according to manufacturer protocol with the low standard procedure.

### Modelling

The mathematical model that we used previously to estimate the contribution of cell-free and cell-to-cell transmission given different environmental conditions has been described in detail within ^7^. In brief, the model describes the turnover and dynamics of (un-)infected CD4+ T cells, CD8+ T cells and the viral load within the different culture systems by ordinary differential equations. The complete set of mathematical equations and detailed description, as well as pre-defined parameter values used within the analyses, are given within^7^. In the original publication, the model was fitted simultaneously to the data of a co-transfer experiment of infected and uninfected CD4+ T cells into 2D suspension, and 3D loose and dense collagen environments, with environmental restriction reducing the infectivity of cell-free virions within collagen, i.e., the transmission parameter 𝛽_𝑓_, to only 14% of the effectivity considered within suspension, i.e., 𝛽_𝑓,𝑙𝑜𝑜𝑠𝑒_ = 𝛽_𝑓,𝑑𝑒𝑛𝑠𝑒_ = 𝜂𝛽_𝑓,𝑠𝑢𝑠_ with 𝜂 = 0.14. Here, we re-performed the analysis done within^7^ by varying 𝜂 between 0 and 1 within steps of 0.1, and also considering 𝜂 = 0.275, as experimentally determined for primary target cells. Fitting was performed as described within^7^ using the *optim*-function within the *R*-language of statistical computing. Posterior distributions of parameter estimates were obtained by performing ensemble fits for each value of 𝜂 based on different starting values. Subsequent filtering steps of fits ensured convergence of parameter estimates by excluding unreasonable dynamics of CD4+ T cell counts and viral loads, with posterior distributions based on ∼110-145 successful fits for each value of 𝜂 (see Sup. Fig 2 b &c).

### TLR inhibitor treatments

To determine the sensing pathway involved in the collagen mediated sensitization of virus particles for innate immune recognition, MDMs were pre-treated with the different inhibitors prior to infection. 5 µM of cGAS inhibitor (G140, Invivogen), 8 µM of TLR 1/2 inhibitor (Cu-CPT22, Selleckchem), 49 µM of TLRs 4& 2/6 inhibitor (GIT27, Tocris), 2 µM of TLR-4 inhibitor (CLI-095, Invivogen), 100 µM of TLR-2 inhibitor (TL2-C29, Invivogen) or 10 µM of TLR-8 inhibitor (Cu-CPT9a, Invivogen) were incubated with MDMs 3h prior to infection. The Myd-88 inhibitor (Pepinh-MYD, Invivogen) was used at 20 µM and incubated with MDMs 4h prior to infection. MDMs were also pre-treated for 1h with a TLR-3/dsRNA complex inhibitor (Merck Millipore) 5 µM final concentration. MDMs were also treated with 100 µM of T20 HIV-1 fusion inhibitor (Roche) or 3 µg/ml Efavirenz (SML0536, Sigma). The different inhibitors were supplemented again during virus inoculation. The cells were then cultured for 3 days at 37°C, and supernatants were harvested for cytokine analysis.

### HIV-1 colocalization with TLR & sub-cellular compartments

MDMs were challenged with NL4.3 R5 Vpr.Int.GFP virions for 4h at 37°C. The plasma membrane of MDMs was then stained using WGA-647 (Invitrogen) for 10 min at 4°C to avoid internalization of the dye, and samples were then fixed using 3% PFA for 90 min. After export out of BSL3 containment, the cells were permeabilized using 0.1% Triton 100X for 10 min at room temperature, incubated for 30 minutes with blocking buffer (PBS, 5% BSA) and then with primary antibodies for 45 minutes at room temperature. After incubation with appropriate secondary antibodies for 30 min at room temperature, the coverslips were mounted onto microscopy slides using Fluoromount-DAPI (Invitrogen). The samples were then imaged using a Leica SP8 point laser scanning confocal microscope using an HC PL APO CS2 63x/1.4 N.A. oil immersion objective. Multichannel images were acquired sequentially using HyD detectors or PMTs in the lightning mode. Z-stacks were acquired with 500 nm steps. Deconvoluted images were then used to segment virions and TLR/compartments using Imaris (Oxford instruments) while enabling object-object statistics. From the resulting data, the percentage of virus segments overlapping with TLR/compartment segments were calculated, using a threshold of 0.01 µm³ overlap to define colocalization.

### Probing HIV-1 Env epitope accessibility

To assess the overall conformation of Env at the surface of virions, we adapted a workflow from (Staropoli et al, 2019). In brief, HIV-1 virions from various strain containing Vpr.Int.GFP fusion proteins were cultured in suspension or in collagen matrices of different densities for 16h. The supernatants were then harvested, and the viral titers were determined by SG-PERT analysis. Equivalent amounts of pURT were then purified by ultracentrifugation over a sucrose cushion using a tabletop ultracentrifuge (TL100, Beckman Coulter). The purified virus pellet was then resuspended in filtered PBS (0.1 µm) and distributed to a 96 well-plate at 1x10^11^ pURT/well. The virions were then incubated with the different antibodies (final concentration: 15µg/ml) or sCD4-PE (1:50 dilution as per manufacturer instructions) for 1 h at 4°C in staining buffer (RMPI with 20% FCS filtered through a 0.1 µm filter). After the first incubation with broadly or non-neutralizing antibodies, the samples were then incubated with a secondary, anti-human IgG1(H+L) AlexaFluor 568 antibody (1:200 dilution; Life Technologies, in staining buffer) for 1h at 4°C. The samples were then fixed using 32% PFA (Thermo Sci) to reach a final concentration of 4%. After export out of BSL-3 containment, the samples were then seeded on poly-L-lysine coated coverslips for 1h, then mounted on microscopy slides. Samples were then imaged using an epifluorescence microscope (Olympus IX81 S1F-3) under a 100X oil objective (PlanApo, N.a. 1.40). Quantification of double positive Vpr-mRuby2 and Alexa Fluor 568 signals was performed using the Spot Detector plugin of the Icy image analysis software. For each primary/secondary antibody pair or sCD4-PE, appropriate signal thresholds were arbitrarily defined to exclude background signals. Background fluorescence thresholds were set using the secondary antibody only condition as a setpoint for unspecific signals. The same thresholds were used to define true positive signals when comparing virions retrieved from suspension or collagen cultures for each antibody.

### Flow cytometry

Samples were measured by flow cytometry in BD FACS Celesta with BD FACS Diva Software. Compensation controls were added for each experiment. Gating was performed using FlowJo software 10.4.2 and data were processed in GraphPad Prism 8.4.3 software.

### Statistical analysis

Statistical analysis of datasets was carried out using Prism version 8.4.3 (GraphPad). For each dataset, normal distribution was tested using the Shapiro-Wilk test, and the appropriate test was used accordingly as indicated in the figure legends. Statistical significance was calculated using one-or two-way ANOVA tests, as well as paired or unpaired t-tests, and Wilcoxon matched paired tests. Correction for multiple comparisons are indicated in figure legends. Significant differences are indicated with the corresponding p-values. Absence of p-values indicates a lack of statistical significance.

## Acknowledgments

This research was funded by the Deutsche Forschungsgemeinschaft (DFG, German Research Foundation) Projektnummer 240245660 – SFB 1129 (project 8 to OTF, project 19 to PC) and 316249678 – SFB 1279 (project A03 to JM). FG was supported by the Chica- and Heinz Schaller Foundation and the German Federal Ministry of Education and Research (BMBF, grant 031L0293A/E). OTF acknowledges support by the Ministry of Science, Research and Culture Baden-Württemberg. This work was also supported by a research grant from the Chica and Heinz Schaller Foundation to PC (Schaller Research Group Leader Programme). We thank the Infectious Diseases Imaging Platform (IDIP) at the CIID at Heidelberg University. We would like to acknowledge access to the infrastructure and support provided by the Cryo-EM Network at Heidelberg University (HDcryoNet), which is funded and supported by the DFG, the Federal Ministry of Science, Research and Culture of Baden-Württemberg, among others, within the framework of the Excellence Strategy of the Federal and State Governments of Germany.

The authors gratefully acknowledge the data storage service SDS@hd supported by the Ministry of Science, Research, and the Arts Baden-Württemberg (MWK), the German Research Foundation (DFG) through grant INST 35/1314-846 1 FUGG and INST 35/1503-1 FUGG. We are grateful to Katharina Morath, Swetha Ananth, Christine Selhuber-Unkel and Ada Cavalcanti-Adam for advice and discussion, to Volker Lohmann, Frank Kirchhoff and Tom Hope for sharing reagents and to Kathrin Bajak for help with preparing the manuscript figures. The following reagents were obtained through the NIH HIV Reagent Program, Division of AIDS, NIAID, NIH: monoclonal anti-Human Immunodeficiency Virus Type 1 (HIV-1) gp120 Protein, Clone 17b (produced *in vitro*), ARP-409, contributed by Dr. James E. Robinson; anti-Human Immunodeficiency Virus (HIV)-1 gp120 Monoclonal Antibody (3BNC117), ARP-12474 and anti-Human Immunodeficiency Virus (HIV)-1 gp120 Monoclonal Antibody (10-1074), ARP-12477 contributed by Dr. Michel Nussenzweig; anti-Human Immunodeficiency Virus (HIV)-1 gp120 Monoclonal Antibody (NIH45-46 G54W), ARP-12174; anti-Human Immunodeficiency Virus (HIV)-1 gp120 Monoclonal Antibody (PG16), ARP-12150, contributed by International Aids Vaccine Initiative.

## Author Contributions

Conceptualization, O.T.F.; Methodology, S.S.A., F.G., Investigation, S.S.A., L.Z., A.I., N.T., K.W.; Data analysis, S.S.A., L.Z.; Mathematical analysis, K.W., F.G.; Writing – Original Draft, O.T.F., S.S.A., L.Z. and F.G.; Writing – Review & Editing, O.T.F, S.S.A., F.G., P.C., J.M.; Funding Acquisition, O.T.F.; Resources, L.R., J.M.; Supervision, O.T.F, P.C., F.G.

---

**Supplementary Figure 1:**
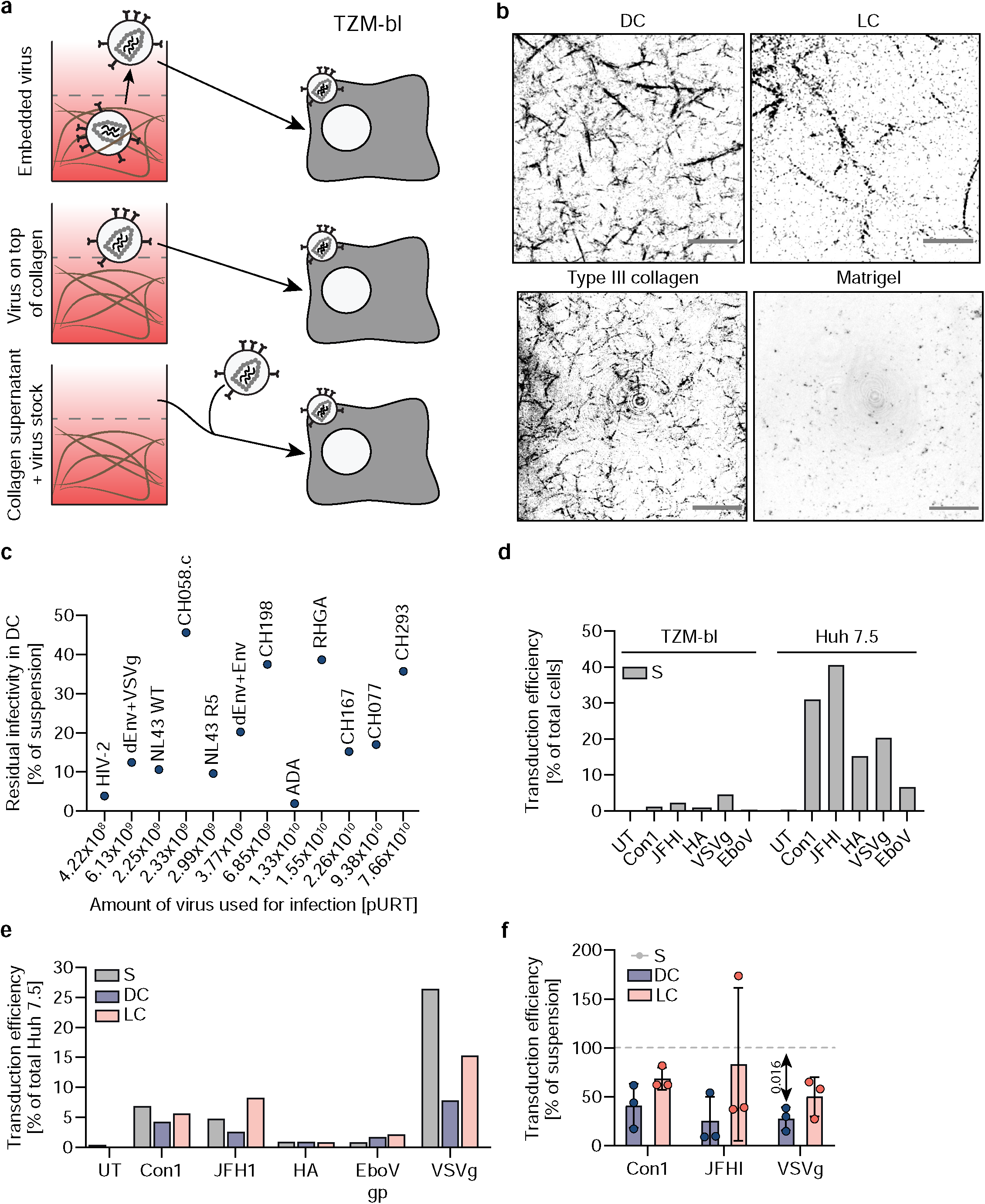
Adhesive matrices exert a restriction on the infectivity of lentiviruses pseudotyped with different viral glycoproteins. **a.** Experimental workflow schematic. Virions were either embedded in collagen matrices, seeded on top of already polymerized collagen gels, or combined with supernatants harvested from collagen matrices polymerized in the absence of virus. The supernatants were then used to infect TZM-bl cells for relative infectivity determination. **b**. Representative micrographs. Different matrices were polymerized and imaged by confocal autoreflection microscopy. **c**. Correlation between the amounts of pURT used for infection for each viral strain and the extent of infectivity restriction after culture in DC gels. **d**. TZM-bl and Huh7.5 cells were transduced with lentiviral particles pseudotyped with different viral glycoproteins and the transduction efficiencies measured by flow cytometry. **e**. More permissive Huh7.5 cells were transduced with the different lentiviral pseudotypes retrieved from suspension or collagen cultures prior to transduction efficiency determination. **f**. Transduction efficiency of Huh7.5 cells with lentiviral particles pseudotyped with the HCV glycoproteins Con1. JFHI or with VSVg after culture in suspension or in collagen. Results represent the mean ± SD from 1 (d,e) or 3 (c,f) independent donors. Significance is indicated by p-values, and was calculated by two-way ANOVA; Tukey’s post-test.

**Supplementary Figure 2:**
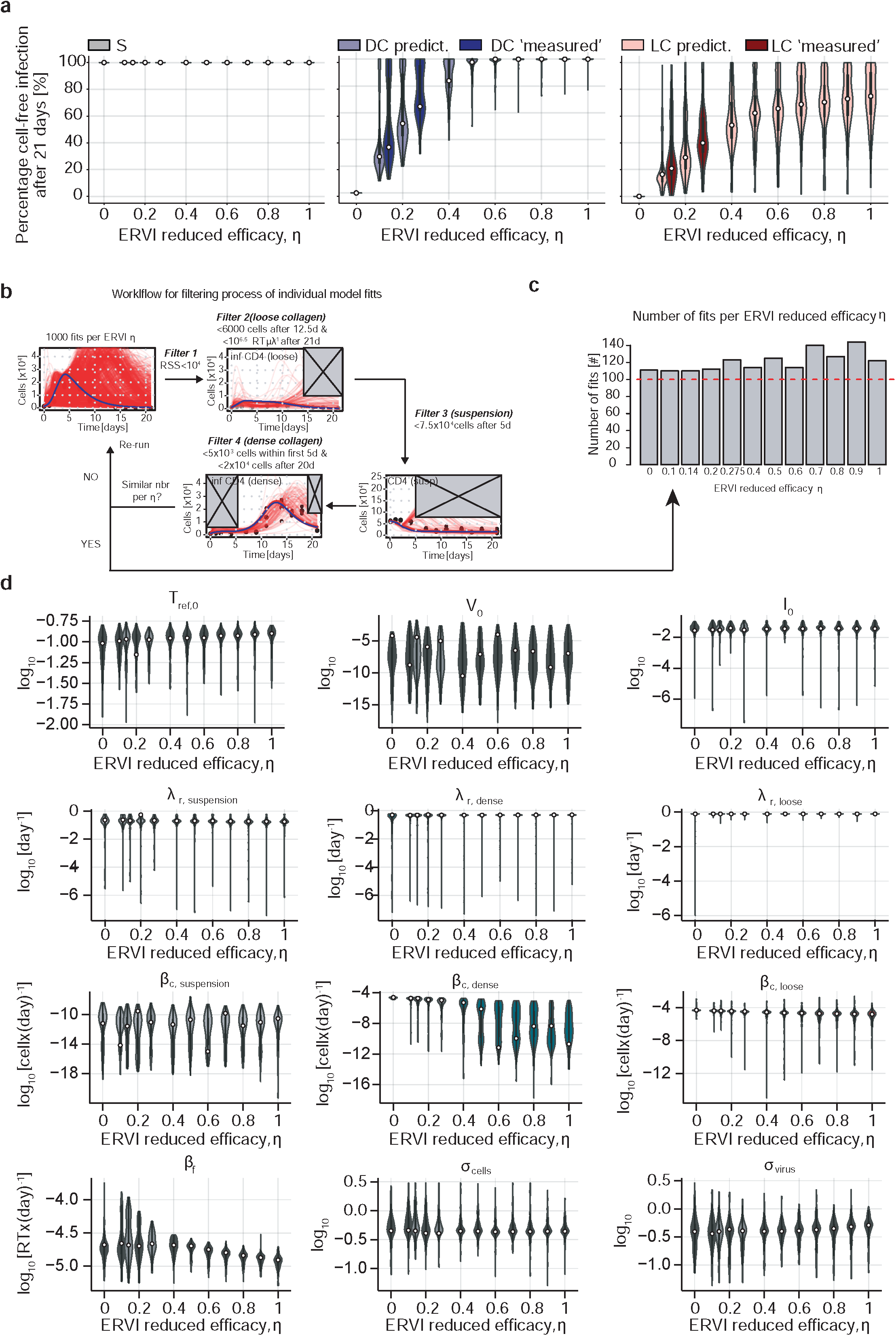
Modelling the consequences of ERVI for HIV-1 spread. **a.** Estimated fraction of cells infected by cell-free transmission after 21 days for the three environmental conditions given different values for the reduced efficacy of virion infectivity by ERVI, 𝜂. The plots show the posterior distributions of estimates over ∼110-140 fits per value of 𝜂 with dots indicating the estimate of the best model fit (see also Materials & Methods). Results for measured values of 𝜂 = 0.14 (Imle et al. 2019) and 𝜂 = 0.275 (here) are shown in dark colours. **b**. Filtering process of individual model fits to ensure comparability of estimates for different reduced efficacies 𝜂, used within the mathematical model shown in (Imle et al. 2019). A sequence of different filtering steps is applied to the results of the ensemble fits given different starting conditions only considering fits with residual sum of squares (RSS) <10^4^ (filter 1), disregarding fits with predictions of >6000 cells after 12.5 days or viral concentrations > 10^6.8^ RT μl^-1^ after 21 days in the supernatant for LC (filter 2), disregarding fits with >7.5×10^4^ CD4 T cells after 5 days in suspension (filter 3), and excluding fits with >5000 CD4 T cells within the first 5 days or >2×10^4^ cells after 20 days for DC. The procedure is repeated to ensure comparable number of estimates for each value of 𝜂 after the filtering for the analysis. **c**. Number of fits per value of 𝜂 considered within the analyses. **d**. Distribution of estimates for the individual parameters describing cell infection dynamics within suspension and 3D collagen environments. Results of the best fit for each value of 𝜂 are indicated by the white dot. For the meaning of the individual parameters and specific model equations see Imle et al., 2019.

**Supplementary Figure 3:**
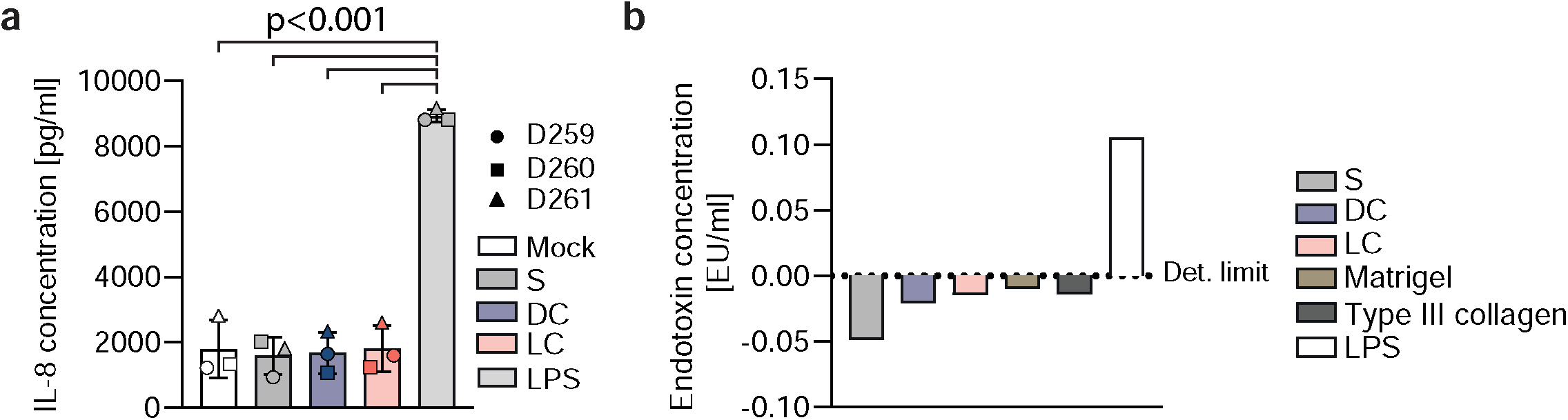
The collagen induced immune sensitization of HIV-1 virions is not caused by soluble components or endotoxins. **a.** Measurement of IL-8 release by ELISA form MDM supernatants. MDMs were cultured with supernatants harvested from suspension or collagen cultures for 72h prior to ELISA analysis. **b**. Endotoxin measurement. The supernatants of suspension, type I or III collagen, or matrigel matrices polymerized in the absence of virus were harvested, and the levels of endotoxin were determined. Results represent the mean ± SD from one experiment (**b**) or 3 independent donors (**a**). Significance is indicated by p-values, and was calculated by one-way ANOVA; Tukey’s post-test.

**Supplementary Figure 4:**
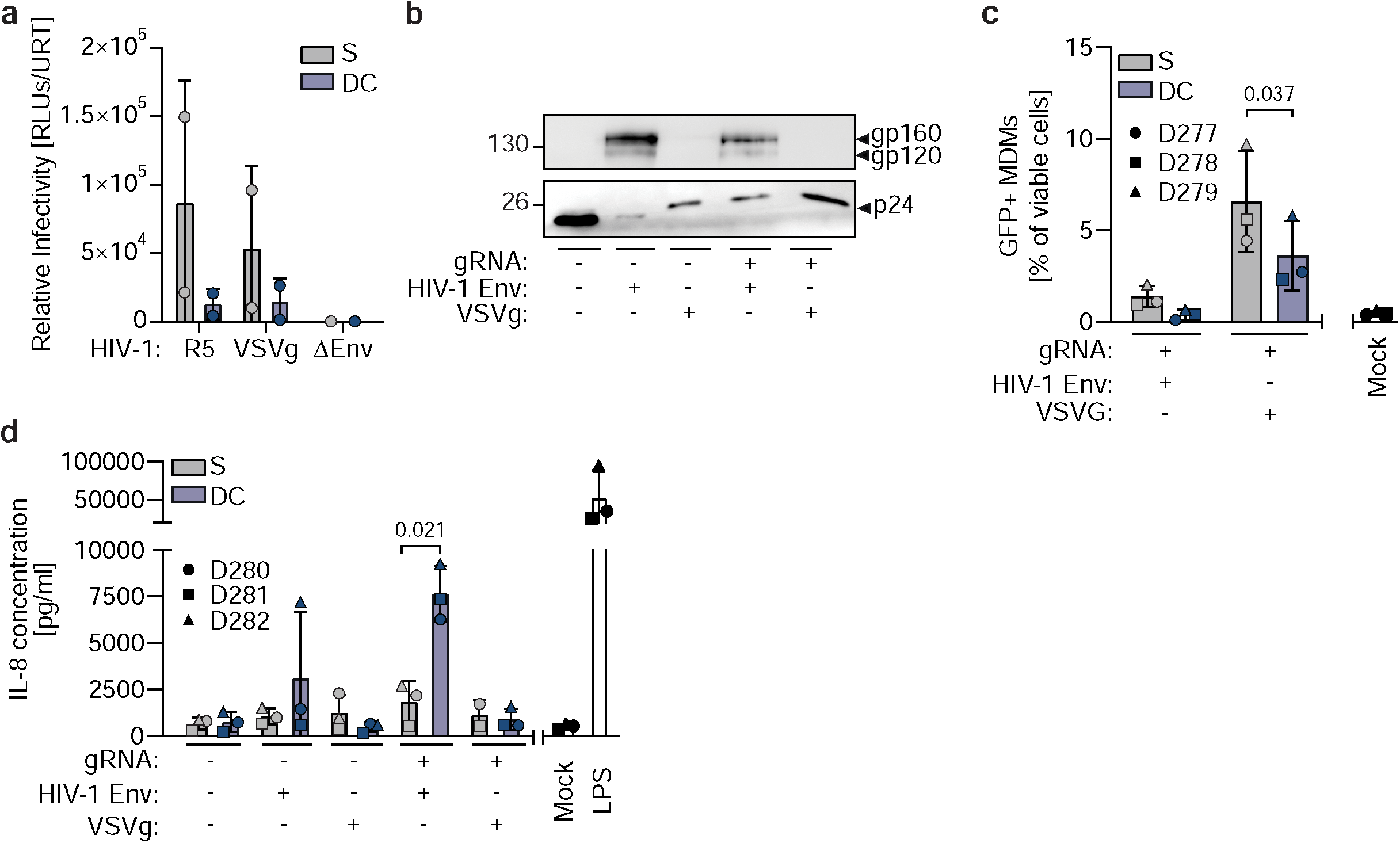
The ERVI induced sensitization depends on the presence of HIV-1 Env and gRNA. **a.** The relative infectivity of HIV-1 NL4.3 R5, HIV-1 ΔEnv VSVg or HIV-1 ΔEnv was determined after 16h of culture in suspension or collagen. **b**. Representative WB result of lentiviruses produced in the presence of HIV-1 Env, VSVg and/or a lentiviral gRNA. Efficient incorporation of p24 and HIV-1 Env was assessed by performing an SDS-PAGE followed by blotting and incubation with anti-p24 or anti-gp120 antibodies. **c**. Flow cytometry analysis of MDMs transduced with lentiviral vectors retrieved from suspension or collagen. Cells that were transduced with gRNA containing lentiviral particles produce GFP, which was detected by flow cytometry. **d**. Measurement of IL-8 release by ELISA from the supernatants of MDMs challenged with the different lentiviral particles. Results represent the mean ± SD from 2 experiments (a) or 3 independent donors (c, d). Significance is indicated by p-values, and was calculated by paired or unpaired t-tests.

**Supplementary Figure 5:**
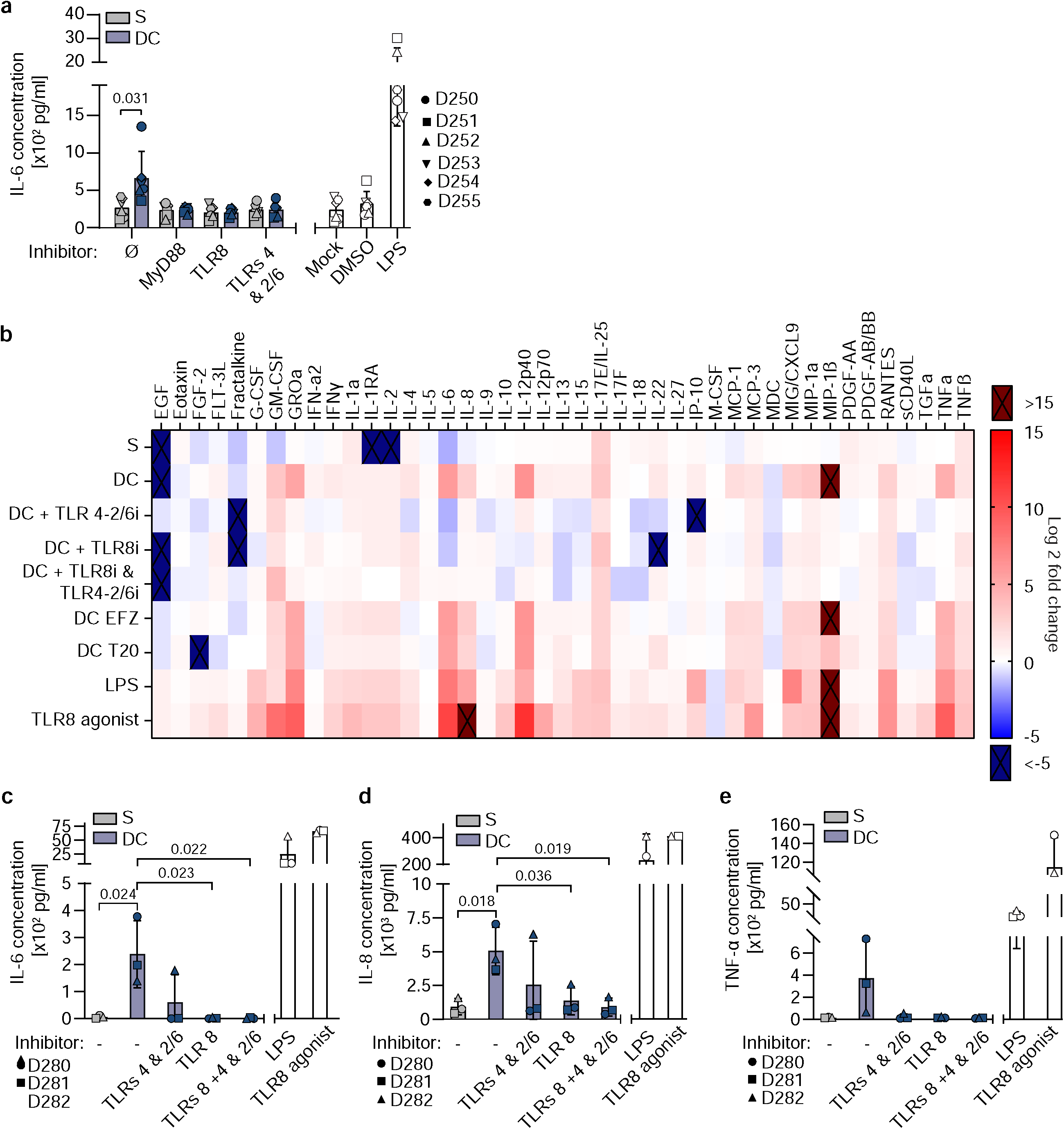
TLRs 2 and 8 recognize collagen experienced particles to elicit a MyD88 dependent innate immune response. **a.** Measurement of IL-6 release by ELISA from the supernatants of MDMs challenged with virions from suspension or collagen cultures in presence or absence of different PRR inhibitors. **b**. Representative heat-map depicting the cytokine profiling for one donor, after challenge with virions from suspension or DC cultures, in presence or absence of HIV or PRR inhibitors. Dark blue & red colors indicate data points below and above the standard curve respectively. **c-e**. Quantification of the levels of IL-6 (**c**), IL-8 (**d**) or TNF-a (**e**) release for 3 donors from the cytokine profiling as in b. Results represent the mean ± SD from 3 (c,d,e) or 6 (a) independent donors. Significance is indicated by p-values, and was calculated by paired or unpaired t-tests.

**Supplementary Figure 6:**
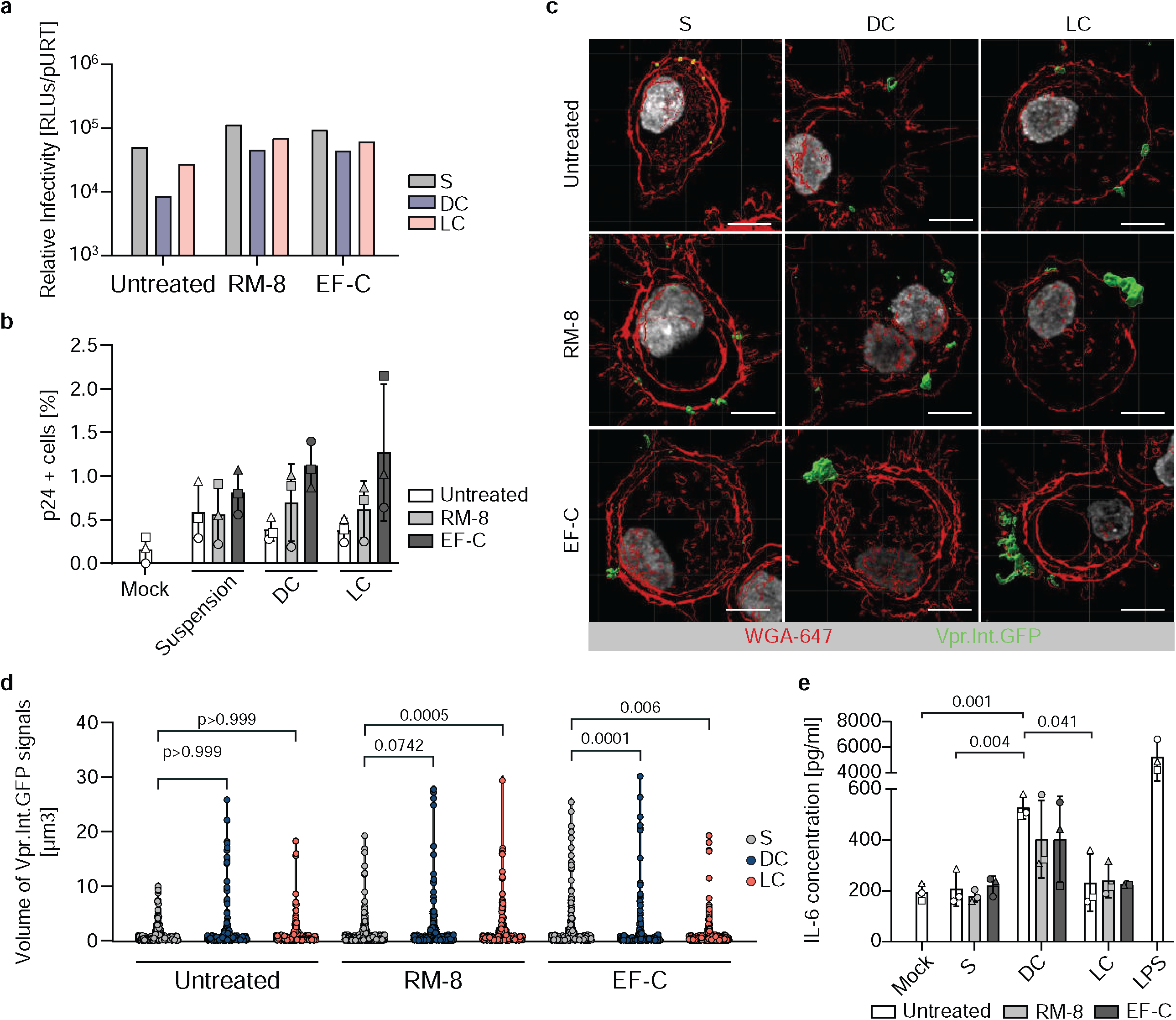
HIV-1 aggregation by infectivity enhancing PNFs does not lead to innate immune sensitization of cell-free virions. **a**. Relative infectivity determination. HIV-1 NL4.3 R5 Vpr.Int.GFP virions were retrieved from suspension or collagen cultures. Equivalent amounts of virions were then incubated with infectivity enhancers and used to infect TZM-bl reporter cells for infectivity determination. **b**. Flow cytometry quantification of the percentage of p24+ MDMs after challenge with virions as in a. **c**. Representative micrographs of MDMs challenged with differentially treated virions. Plasma membrane segmentation: red, virus segmentation: green. **d**. Quantification of the volumes of segmented virions as in c. **e**. Measurement of the levels of IL-6 release by ELISA from the supernatants of MDMs challenged with differentially treated virions. Results represent the mean ± SD from 1 experiment (a) or 3 independent donors (b,d,e). Significance is indicated by p-values, and was calculated by two-way ANOVA, Tukey’s post-test (b); Kruskal-Wallis test, Dunn’s post-test (d) or one-way ANOVA, Tukey’s post-test (e).

**Supplementary Figure 7:**
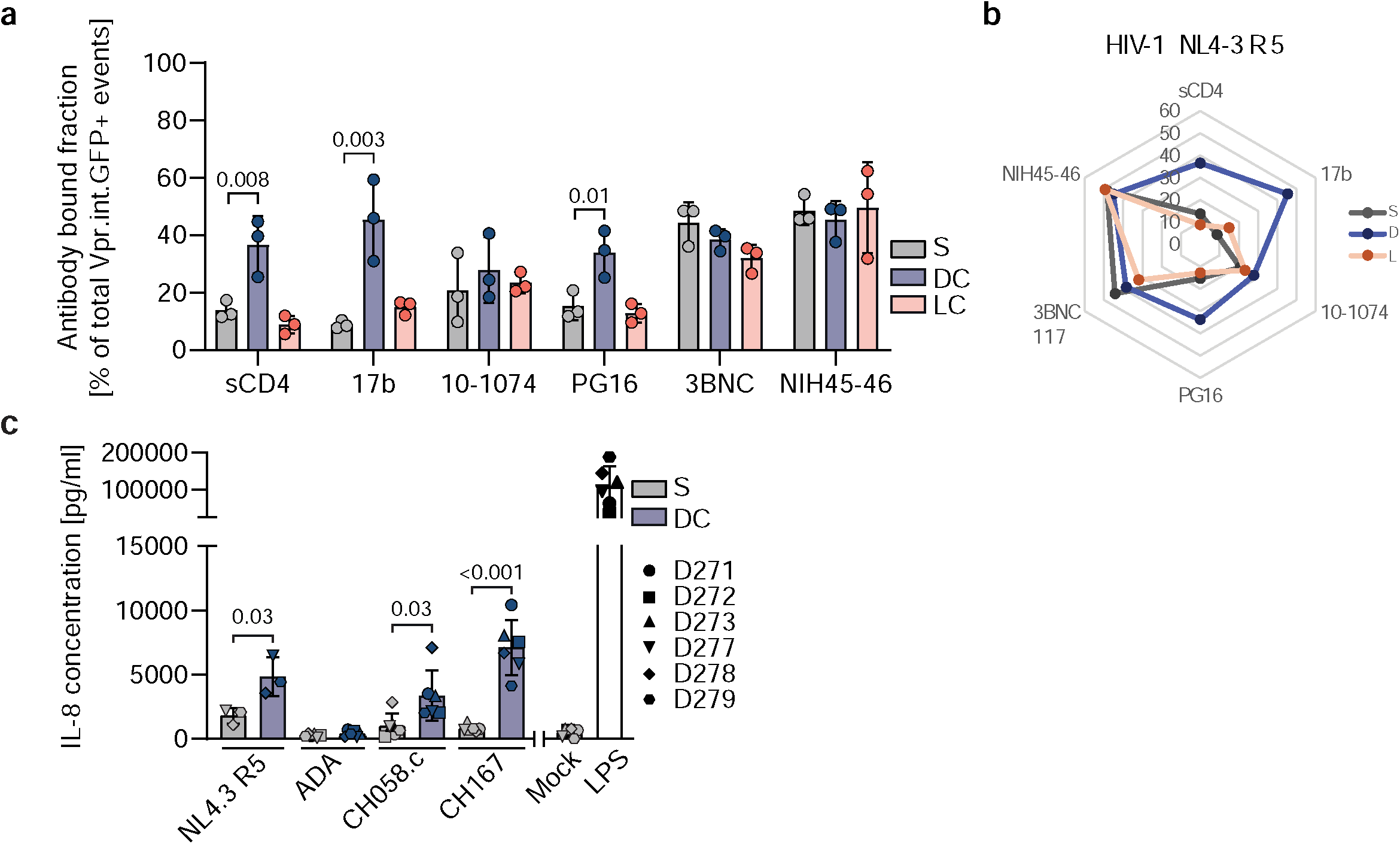
HIV-1 Env is structurally rearranged after contact of virions with collagen fibers. **a**. Quantification of the frequency of antibody/sCD4 binding to differentially cultured HIV-1 NL4.3 R5 Vpr.Int.GFP virions as shown in Fig. 7c. **b**. Spider plot representing the frequency of antibody binding to HIV-1 NL4.3 R5 as in (a). **c**. Measurement of IL-8 release by ELISA from the supernatants of MDMs challenged with different HIV-1 strains subjected to suspension or collagen cultures. Results represent the mean ± SD from 3 independent donors. Significance is indicated by p-values, and was calculated by paired t-tests.

**Supplementary Figure 8:**
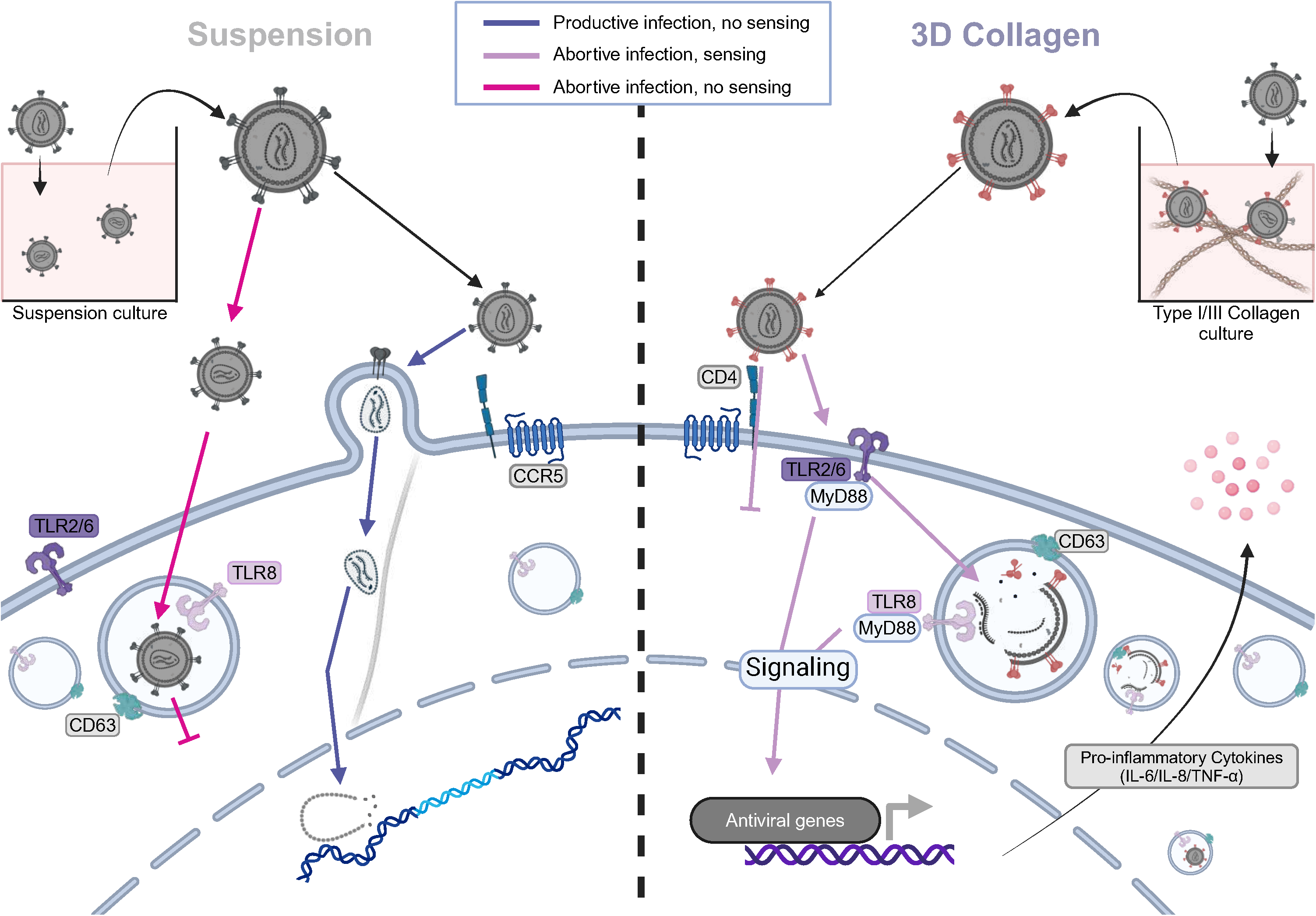
Schematic model of the dual mechanisms by which ERVI impairs the infectivity of HIV-1 particles and sensitizes them for TLR-2 and TLR-8 innate immune recognition in MDMs. See discussion for details.

**Supplementary Movie 1: Localization to TLR8 containing endosomes of virions derived from suspension cultures.**

**Supplementary Movie 2: Localization to TLR8 containing endosomes of virions derived from dense collagen cultures.**

